# Unified fate mapping in multiview single-cell data

**DOI:** 10.1101/2023.07.19.549685

**Authors:** Philipp Weiler, Marius Lange, Michal Klein, Dana Pe’er, Fabian J. Theis

## Abstract

Single-cell RNA sequencing allows us to model cellular state dynamics and fate decisions using expression similarity or RNA velocity to reconstruct state-change trajectories. However, trajectory inference does not incorporate valuable time point information or utilize additional modalities, while methods that address these different data views cannot be combined and do not scale. Here, we present CellRank 2, a versatile and scalable framework to study cellular fate using multiview single-cell data of up to millions of cells in a unified fashion. CellRank 2 consistently recovers terminal states and fate probabilities across data modalities in human hematopoiesis and mouse endodermal development. Our framework also allows combining transitions within and across experimental time points, a feature we use to recover genes promoting medullary thymic epithelial cell formation during pharyngeal endoderm development. Moreover, we enable estimating cell-specific transcription and degradation rates from metabolic labeling data, which we apply to an intestinal organoid system to delineate differentiation trajectories and pinpoint regulatory strategies.

## Introduction

Single-cell assays uncover cellular heterogeneity at unprecedented resolution and scale, allowing complex differentiation trajectories to be reconstructed using computational approaches^1–10^. While these trajectory inference (TI) methods have uncovered numerous biological insights^9, 11^, they are typically designed for snapshot single-cell RNA sequencing (scRNA-seq) data and cannot accommodate additional information relevant for understanding cell-state dynamics, including experimental time points, multimodal measurements, RNA velocity^12, 13^, and metabolic labeling^14–21^.

We and others have developed methods to analyze emerging data modalities, such as CellRank^22^ for RNA velocity, Waddington-OT^23^ for experimental time points, and dynamo^24^ for metabolic labeling data. However, each method only addresses a single modality, thereby ignoring much of the upcoming multi-modal information for trajectory analysis. This specialization renders many biological systems inaccessible; for example, adult hematopoiesis violates assumptions of current RNA velocity models^13, 22^, precluding us from applying CellRank to this well-studied system and prompting the question of whether the algorithm could be developed further to reconstruct differentiation dynamics using another aspect of this data.

To address this challenge, we decompose trajectory inference into two components - modality-specific modeling of cell transitions, followed by modality-agnostic trajectory inference - and developed CellRank 2, a robust, modular framework to analyze multiview data from millions of cells. CellRank 2 generalizes CellRank and is capable of exploiting the full potential of new data modalities to study complex cellular state changes and identify initial and terminal states, fate probabilities, and driver genes.

We demonstrate CellRank 2’s flexibility across a series of applications: Using a pseudotime in a hematopoiesis context, determining developmental potentials during embryoid body formation, and combining experimental time point with intra-time point information for pharyngeal endoderm development. Our approach for incorporating experimental time recovers terminal states more faithfully, allows studying gene expression change continuously across time, and discovers putative progenitors missed by alternative approaches. We also introduce a new computational approach for learning cellular dynamics from metabolic labeling data and show that it reveals regulatory mechanisms by recovering cell-specific transcription and degradation rates in mouse intestinal organoids.

## Results

### A modular framework for studying state-change trajectories

CellRank 2 models cell-state dynamics from multiview single-cell data. It automatically determines initial and terminal states, computes fate probabilities, charts trajectory-specific gene expression trends, and identifies putative driver genes (Methods). A broad and extensible range of biological scenarios can be studied using its robust, scalable, and modular design (Methods).

Similar to CellRank^22^, we employ a probabilistic system description wherein each cell constitutes one state in a Markov chain with edges representing cell-cell transition probabilities. However, we now enable deriving these transition probabilities from various biological priors. Following previous successful trajectory inference approaches^1–9^, we assume gradual, memoryless transitions of cells along the phenotypic manifold. The assumption of memoryless transitions is justified as we model average cellular behavior (Methods).

To allow broad applicability, we divide CellRank 2 into *kernels* for computing a cell-cell transition matrix based on multiview single-cell data and *estimators* for analyzing the transition matrix to identify initial and terminal states, compute fate probabilities, and perform other downstream tasks. CellRank 2 provides a set of diverse kernels that derive transition probabilities based on gene expression, RNA velocity, pseudotime, developmental potentials, experimental time points, and metabolically labeled data (Fig. 1a and Supplementary Fig. 1). For an initial, qualitative overview of recovered cellular dynamics, we introduce a random walk-based visualization scheme (Methods).

**Fig. 1.**
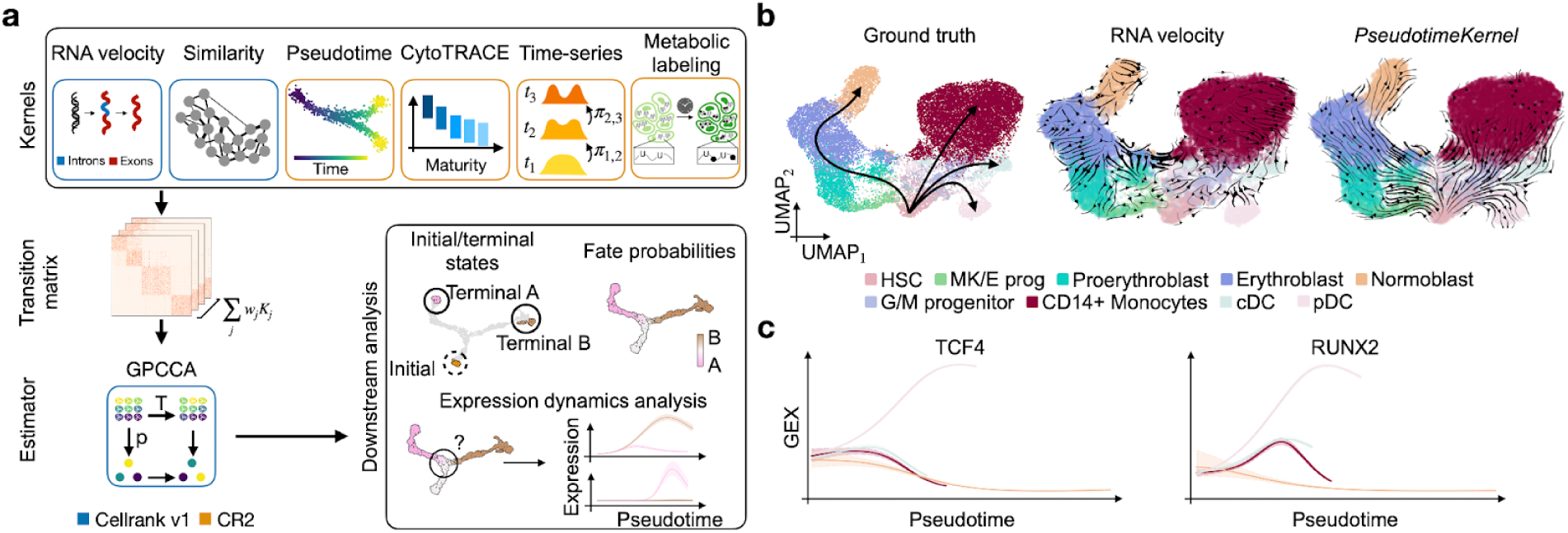
| CellRank 2 provides a unified framework for studying single-cell fate decisions using Markov chains. **a.** CellRank 2 (CR2) uses a modular design. Data and problem-specific *kernels* calculate a cell-cell transition matrix inducing a Markov chain (MC). *Estimators* analyze the MC to infer initial and terminal states, fate probabilities, and driver genes. Using fate probabilities and a pseudotime allows for studying gene expression during lineage priming. Features inherited from the original CellRank implementation are indicated in blue, new features in orange. **b.** UMAP embedding of 24,440 peripheral blood mononuclear cells (PBMCs)^31^, colored by cell type (HSC: hematopoietic stem cell, MK/E prog: megakaryocyte/erythrocyte progenitors, G/M progenitor: Granulocyte/Myeloid progenitor, pDC: plasmacytoid dendritic cell, cDC2: classical dendritic cells). The well-known differentiation hierarchy (left) and projected velocity fields based on RNA velocity (middle) and the *PseudotimeKernel* (right) are colored in black. **c.** Correlating fate probabilities with gene expression recovers lineage drivers for the pDC lineage. For each lineage, the gene expression trend is weighted by the fate probability; color coding corresponds to lineage.

For many biological processes, the starting point can be quantified robustly and cells ordered along a pseudotime. We propose to use this fact by biasing the edges of a nearest neighbor graph towards mature cell states to estimate cell-cell transitions. Developmental potentials can be used similarly. CellRank 2 generalizes earlier concepts^7, 25^ to arbitrary pseudotimes and atlas-scale datasets with the *PseudotimeKernel* and *CytoTRACEKernel*. More complex systems with unknown initial states or longer developmental time scales can be captured faithfully through multiple experimental time points. To reconstruct the overall developmental dynamics described both across and within time points, we extend classical optimal transport^23^, in particular WaddingtonOT^23^, with our *RealTimeKernel* to include within time point dynamics. In contrast, metabolic labels offer an experimental approach to overcome the discrete nature of distinct experimental time points^14, 15, 18^. Based on this information, we developed an inference approach quantifying kinetic rates that allow us to infer cell transitions. In the following, we provide details on each kernel and demonstrate the versatility of our approach through diverse applications. Finally, various kernels may be combined to yield a more complete picture of cellular dynamics through multiview modeling.

Once we have inferred a transition matrix, we use an estimator module^26, 27^ to uncover cellular trajectories, including initial and terminal states, fate probabilities, and driver genes. Critically, estimators are view-independent and are, thus, applicable to any transition matrix (Methods). To scale these computations to large datasets, we assume that each cell gives rise to only a small set of potential descendants. This assumption yields sparse transition matrices for every kernel and allows CellRank 2 to compute fate probabilities 30 times faster than CellRank (Supplementary Fig. 2a; Methods).

The modular and robust design makes CellRank 2 a flexible framework for the probabilistic analysis of state dynamics in multiview single-cell data: It enables the rapid adaptation of computational workflows to emerging data modalities, including lineage tracing^28^ and spatiotemporal data^29^, the support of new data modalities with kernels, and the support of new analyses with estimators (Methods).

### Overcoming RNA velocity limitations

RNA velocity infers incorrect dynamics in steady-state human hematopoiesis due to violated model assumptions, even though pseudotime can be recovered faithfully (Supplementary Fig. 3; Supplementary Note 1). Specifically, the assumption of constant rates made by conventional RNA velocity models is violated, and genes important for this system exhibit high noise or low coverage. The remarkable success of traditional pseudotime approaches in systems with well-known initial conditions motivated us to circumvent RNA velocity limitations by developing the *PseudotimeKernel*, which computes pseudotime-informed transition probabilities (Methods) and a corresponding vector field. Building upon conceptual ideas proposed for Palantir^7^, our approach generalizes to any precomputed pseudotime and uses a soft weighting scheme^30^.

We applied the *PseudotimeKernel* to human hematopoiesis^31^ and computed transition probabilities based on diffusion pseudotime (DPT)^2^ for the normoblast, monocyte, and dendritic cell lineages (Fig. 1b and Supplementary Fig. 4a). The *PseudotimeKernel* correctly recovered all four terminal states with fate probabilities reproducing ground truth (Supplementary Fig. 4b,c). To additionally visualize the recovered dynamics, we generalized the streamline projection scheme from RNA velocity^12, 13^ to any neighbor-graph-based kernel (Fig. 1b). We identified putative gene candidates driving lineage commitment by correlating gene expression with lineage-specific fate probabilities (Methods). This approach correctly identified the transcription factors RUNX2 and TCF4 as regulators of the pDC lineage^32, 33^ (Fig. 1c).

Compared to the *PseudotimeKernel*, an RNA velocity-based analysis failed to recover the cDC lineage (Supplementary Fig. 4b), and fate probabilities assigned by the *VelocityKernel* violated the known lineage commitment and hierarchy, including high transition probabilities from proerythroblast and erythroblast cells to monocytes instead of normoblasts (Supplementary Fig. 4c).

Our *PseudotimeKernel* generalizes to any pseudotime, allowing users to choose the algorithm most suitable for their dataset^34^. In systems with simpler differentiation hierarchies and known initial states, CellRank 2’s *PseudotimeKernel* yields additional insights into terminal states and fate commitment compared to classical pseudotime approaches.

### Learning vector fields from developmental potentials

Pseudotime inference requires the initial state to be specified. The *PseudotimeKernel* is, thus, only suitable for systems with known differentiation hierarchies. If the initial state is not known, CytoTRACE^25^ can be used to infer a stemness score by assuming that, on average, naive cells express more genes than mature cells. We found this assumption to be effective for many early developmental scenarios, but, critically, CytoTRACE does not scale in time and memory usage when applied to large datasets and does not resolve individual trajectories through terminal states and fate probabilities (Supplementary Fig. 2b). We thus developed the *CytoTRACEKernel* by revising the original CytoTRACE approach, such that edges of k-nearest neighbor graphs point toward increasing maturity and quantify cell-cell transition probabilities on atlas-scale datasets (Supplementary Fig. 5a; Methods). Results from our kernel agree with the original approach across multiple datasets (Supplementary Fig. 5b,c; Methods). Further, we compared computational performance on a mouse organogenesis atlas^8^ containing 1.3 million cells. While the original implementation failed above 80,000 cells, our adaptation ran on the full dataset in under two minutes (Supplementary Fig. 2b).

We applied the *CytoTRACEKernel* to study endoderm development in pluripotent cell aggregates known as embryoid bodies^35^ (Supplementary Fig. 6a). The CytoTRACE-based pseudotime increased smoothly throughout all experimental time points, as expected (Supplementary Fig. 6b), and allowed us to identify 10 of 11 terminal cell populations (Supplementary Fig. 6c). In contrast, Palantir and DPT identified a bimodal population distribution of early cells in the first stage, resulting in a compressed range of pseudotimes for all other populations and stages (Supplementary Fig. 6b).

The endoderm gives rise to internal organs; thus, we correlated fate probabilities with gene expression to infer putative driver genes which may direct organogenesis, identifying the MIXL1, FOXA2, and SOX17 transcription factors (TFs), in agreement with the original publication^35^ (Supplementary Fig. 6d). To uncover potential upstream regulators of these TFs, we visualize smooth gene expression trends of top-ranked putative drivers of the endoderm trajectory in a heatmap and sorted genes according to their peak in CytoTRACE pseudotime (Supplementary Fig. 6e). We found LINC00458, LINC01356, NODAL, IFITM1, and 9 TFs to peak before FOXA2 and SOX17. All are known mouse endodermal development genes^36–38^, and our prediction that LINC00458 expression peaks before LINC01356 has been observed previously^36^ as well.

CellRank 2’s *CytoTRACEKernel* allowed us to infer cellular dynamics from a snapshot of endoderm development without having to specify an initial state for pseudotime computation. We recovered terminal states, known driver genes, and their temporal activation pattern.

### Adding a temporal resolution to fate mapping

Single-cell time series are increasingly popular for studying non-steady-state differentiation programs. The computational challenge lies in matching cells sequenced at different time points to reconstruct trajectories of state change. Previous methods have either determined a general pseudotime^39^ or used optimal transport (OT) for this task^23^ but ignore transitions within time points that contain valuable information for directing transitions and detecting terminal states. We developed the *RealTimeKernel,* which combines Waddington-OT (WOT)-computed^23^ inter-time-point transitions with similarity-based intra-time-point transitions to allow for multiview modeling (Fig. 2a; Methods). Importantly, considering inter-time point transitions enables unbiased identification of terminal states from time-course studies through a more granular cell fate mapping (Fig. 2b and Supplementary Fig. 7a,b).

**Fig. 2.**
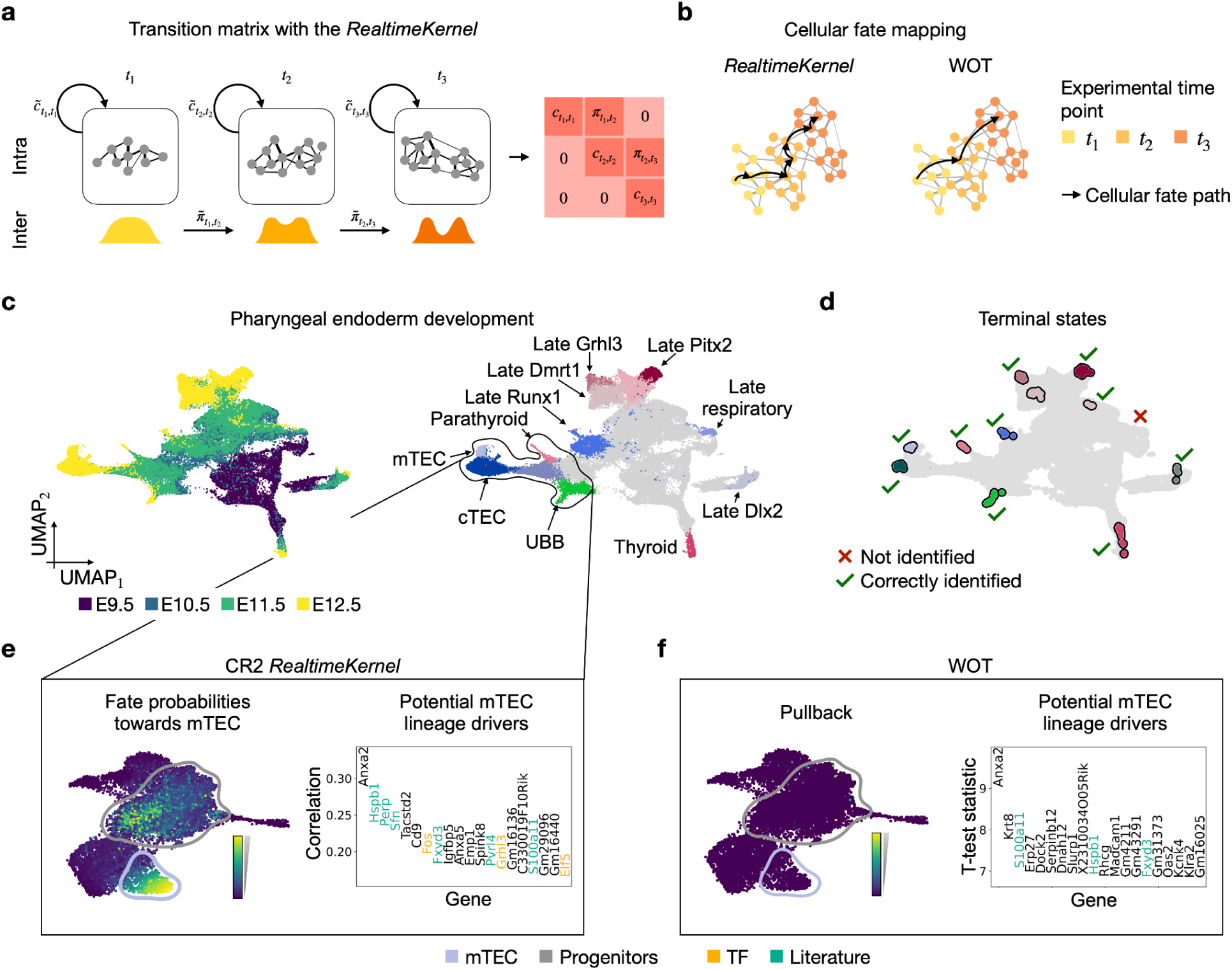
| Inferring state trajectories through time-resolved measurements. **a.** The *RealTimeKernel* combines across time point transitions from WOT^23^ with within time point transitions from gene expression similarity to account for the asynchrony observed in many cellular processes. All views are combined in a single transition matrix. **b.** By including within-time point information, the *RealTimeKernel* enables recovering more granular state transitions. WOT only considers transitions between consecutive time points. **c.** UMAP embedding of pharyngeal organ development^40^ (N = 55,044 cells) colored by embryonic day (E; left) and original cell type annotation (right, mTEC: medullary thymic epithelial cell, cTEC: cortical thymic epithelial cell, UBB: ultimobranchial body). Grey color encodes early, uncommitted cells. **d.** Using the *RealTimeKernel*, CellRank 2 correctly identifies 10 out of 11 terminal states. The black outline highlights mTECs and potential precursor cells. **e,f.** Fate probabilities towards the mTEC terminal state (left) and top 20 putative driver genes identified (right) based on the *RealTimeKernel* (**e**) or WOT’s pullback distribution (**f**). We highlight TFs in yellow and known mTEC development genes in green. CellRank 2 identifies putative drivers by correlating fate probabilities with gene expression, WOT by comparing high- and low-probability cells.

Many OT implementations, including WOT^23^, use entropic regularization^41^ to speed up computation. However, this practice introduces dense transition matrices, which slows downstream applications, hindering us from analyzing large datasets. We, therefore, developed an adaptive thresholding scheme to sparsify transition matrices (Methods), yielding 9-fold and 56-fold faster macrostate and fate probability computation, respectively, on a mouse embryonic fibroblast (MEF) reprogramming dataset^23^ (Supplementary Fig. 2c). To validate our thresholding scheme, we correlated fate probabilities towards the four terminal states, with and without thresholding, and found very high correlations within each lineage (Pearson correlation coefficient ; Supplementary Fig. 7c; Methods).

Numerous applications, including gene trend plotting and driver gene identification, require continuous temporal information rather than discrete time points. Thus, we comprised a new, real-time informed pseudotime approach, which uses experimental time points but embeds them in the continuous landscape of expression changes (Methods). As expected in this system, our new pseudotime indeed correlates better with experimental time than traditional pseudotime approaches on the MEF reprogramming data (Supplementary Fig. 8b,c). Compared to WOT, we enable studying gradual fate establishment along a continuous axis (Supplementary Fig. 8d-g).

The pharyngeal endoderm, an embryonic tissue, plays a crucial role in patterning the pharyngeal region and developing organs^40^, such as the parathyroid, thyroid, and thymus^42–44^. Multiple experimental time points can capture its development, making it an ideal candidate system for our *RealTimeKernel*. We analyzed gene expression change from embryonic days (E) 9.5 to 12.5^40^ (Fig. 2b) and automatically recovered ten of the eleven terminal states manually assigned in the original publication (Fig. 2c). Correlating fate probabilities with gene expression correctly recovered known lineage drivers of the parathyroid (*Gcm2*)^45^, thyroid (*Hhex*)^46^, and thymus (*Foxn1*)^47^ (Supplementary Fig. 9).

To disentangle the trajectory leading to medullary thymic epithelial cells, a stromal cell type associated with thymic adhesion^48, 49^, we first took the subset of parathyroid, ultimobranchial body, medullary and cortical thymic epithelial cells (mTECs, cTECs), and their progenitors (Supplementary Fig. 10a; Methods). Computing fate probabilities towards terminal states, we discovered a progenitor cell cluster with an increased probability of assuming mTEC fate (Fig. 2d and Supplementary Fig. 10b). It is easy to overlook this putative mTEC ancestor cluster in the 2D UMAP embedding, highlighting the importance of our high-dimensional fate analysis (Supplementary Fig. 10c). Next, we used our correlation-based analysis to identify possible drivers of this fate decision and found genes relevant for the thymus lineage (*Hspb1*, *Perp*, *Sfn*, *Fxyd3*, *Pvrl4*, *S100a11*)^50–54^ as well as TFs (*Fos*, *Grhl3*, *Elf5*) among the 20 genes with highest correlation (Fig. 2d).

Unlike CellRank 2, WOT relies solely on inter-time-point information. Applied to the pharyngeal endoderm data, it failed to identify the putative mTEC ancestor cluster. Additionally, even when we leveraged the knowledge of putative mTEC progenitors identified by the *RealTimeKernel* at the earlier experimental time points, classical WOT identified fewer driver gene candidates with known functions in mTEC development at these time points (Fig. 2e and Supplementary Fig. 10d,e). We speculate that this decrease in performance is caused by WOT relying on pullback distributions, which assign a likelihood to each early-day cell to differentiate into any late-day cell but do not take intra-time point dynamics into account. In contrast, CellRank 2 computes continuous fate probabilities with a global transition matrix, combining transitions within and across time points (Methods). Finally, classical differential expression testing also recovered fewer known driver genes and TFs as our correlation-based analysis (Supplementary Fig. 10f).

Our *RealTimeKernel* incorporates gene expression changes within and across experimental time points. Notably, these complementary views allowed identifying a putative progenitor population and substantially more relevant drivers compared to approaches focusing on a single data view.

### Estimating cell-specific kinetic rates and fate with metabolic labeling data

The destructive nature of standard single-cell protocols prohibits directly examining gene expression changes over time. Metabolic labeling of newly transcribed mRNA molecules^14, 15, 17, 18^, however, yields time-resolved single-cell RNA measurements^55^ that should substantially improve our ability to learn system dynamics. The temporal resolution is in the order of minutes to hours and thus much finer compared to typical time course studies. We developed an approach to learn directed state-change trajectories from metabolic labeling data using pulse- and chase experiments (Fig. 3a; Methods).

**Fig. 3.**
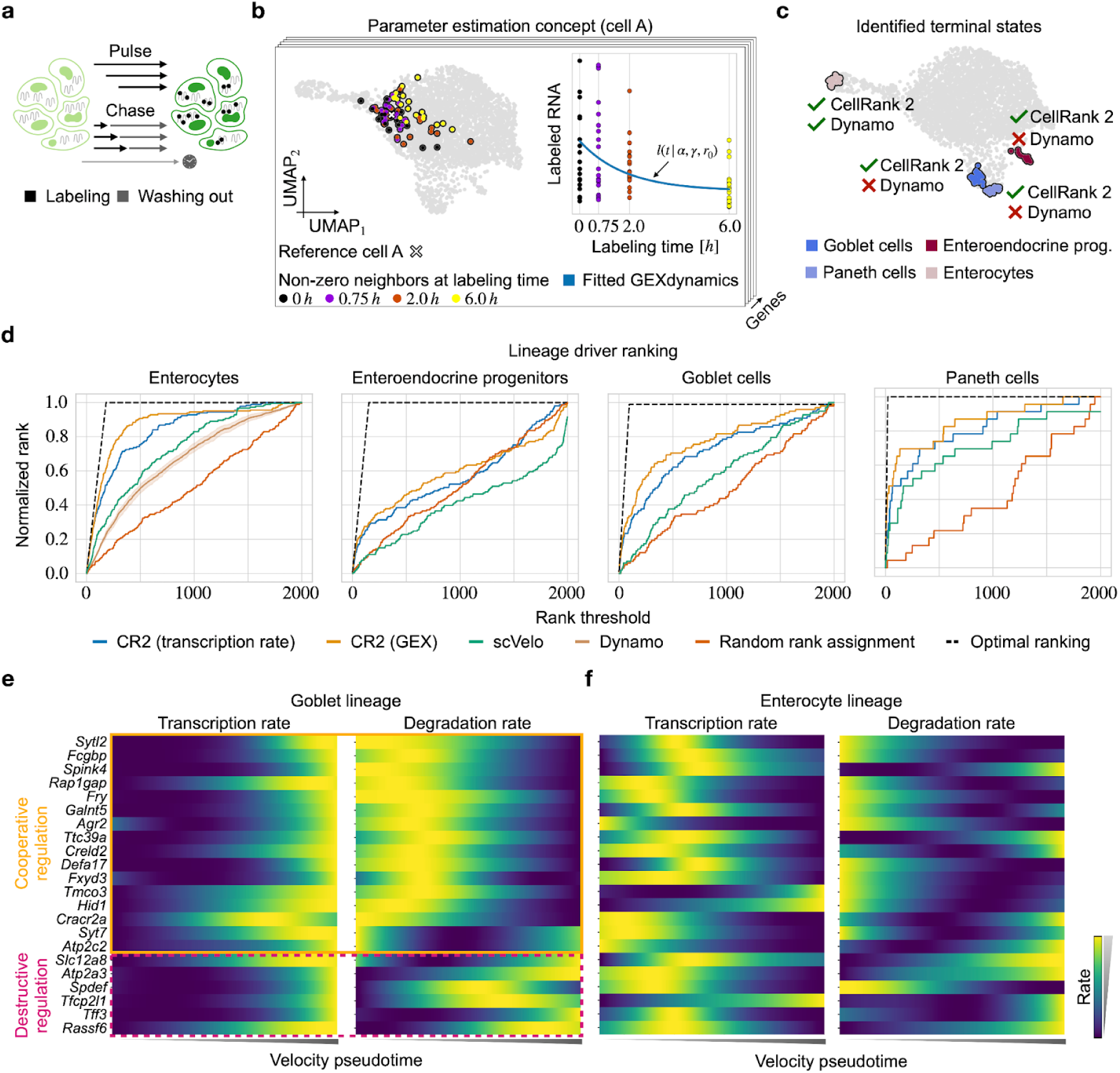
| Quantifying lineage-specific regulation strategies through metabolic labeling. **a.** Cells are metabolically labeled in pulse experiments^15^, involving incubation with nucleoside analogues for varying durations followed by simultaneous sequencing. In chase experiments, in which cellular mRNA fully incorporates nucleoside analogues during a long incubation, then washing out of these nucleosides for varying durations, followed by simultaneous sequencing of all cells. **b.** For each cell, gene, and labeling duration, we identify the number of neighbors such that a pre-defined number of cells with non-trivial counts are included in the neighborhood, illustrated here for an exemplary gene A. These cells are then used to estimate cell and gene-specific transcription and degradation rates and, respectively, to model the dynamics of labeled mRNA. **c.** UMAP embedding highlighting terminal states identified using CellRank 2 and dynamo, respectively. **d.** Ranking of drivers for each lineage identified by different methods (CR2: CellRank 2). Dynamo identified only enterocytes as terminal and, thus, provides a gene ranking only for this lineage. For dynamo, the mean gene ranking and corresponding 95% confidence band are shown (Methods). **e.** Inferred transcription (left) and degradation (right) rates of top-ranked known drivers of the goblet lineage. **f.** Same as **e** but along the enterocyte lineage.

Similar to previous approaches^15^, we model mRNA dynamics through a dynamical system, including mRNA molecule transcription and degradation rates^15^. We estimate these rates for each cell- and gene by considering the dynamical information conveyed through metabolic labels (Fig. 3b; Methods). Based on a cell-cell similarity graph, for each cell, gene, and labeling time, we identify a neighborhood in which a sufficient number of cells express the given gene. Next, we estimate transcription and degradation rates based on these cell sets by minimizing the squared distance between observed and estimated transcripts. Based on these parameters, we infer a high-dimensional velocity vector field used to obtain cell-cell transition probabilities with the *VelocityKernel*.

We applied our devised method to data from murine intestinal organoids labeled with scEU-seq^15^, focusing on the enterocyte, enteroendocrine, goblet, and paneth lineages (Supplementary Fig. 11a; Methods). Following parameter estimation, we computed the underlying velocity field, inferred transition probabilities, and recovered all four terminal states (Fig. 3c). We assessed the quality of inferred terminal states via cell-type purity, defined as the percentage of the most abundant cell type, reasoning that a high cell-type purity results from a low inference uncertainty of the underlying transition matrix (Methods). Indeed, we observed a high cell-type purity (85% on average) for each terminal state (Supplementary Fig. 11b).

We compared our approach with dynamo^24^, an alternative method for estimating cellular dynamics based on metabolic labeling data. In contrast to our approach, dynamo relies on a steady-state assumption, only uses a small subset of cells for parameter inference, does not estimate cell-specific rates, and infers cellular trajectories deterministically. Applied to the organoid data, dynamo only recovered the enterocyte population as a terminal state (Fig. 3c and Supplementary Fig. 11c; Methods).

Beyond identifying the most mature cell population in each lineage, we asked whether our approach ranked known lineage drivers higher than competing approaches that do incorporate labeling information (dynamo) or that do not (scVelo’s *dynamical model* of RNA velocity and a random baseline). To assess the quality of each method’s gene ranking, we curated an optimally ranked list of known regulators and markers^56^ of each lineage and compared each method’s ranking to it (Methods). As dynamo only identified enterocytes as a terminal cluster, it could not rank drivers of any other lineage. Our approach achieved the best ranking for each of the four terminal states (Fig. 3d and Supplementary Fig. 11d) and notably outperformed competing approaches both when correlating gene expression and the inferred transcription rates with fate probabilities to identify driver genes.

The estimated cell- and gene-specific kinetic rates enabled us to investigate how these drivers are regulated by mRNA transcription and degradation. Analyzing the regulatory strategies of known markers and regulators ranked among the top 100 putative drivers for the goblet lineage revealed two different regulatory strategies (Fig. 3e). The first strategy increases transcription rates with decreasing degradation rates (*e.g.*, *Spdef*, *Sytl2*, *Fcgbp*), and the second simultaneously increases transcription and degradation rates (*e.g.*, *Atp2a3*, *Tff3*, *Rassf6*); both align with earlier findings of cooperative (case 1) and destructive (case 2) regulation strategies^15^. Similarly, in the enterocyte lineage, this same set of genes predominantly exhibits either (1) a decreased transcription accompanied by an increase in degradation rate (cooperative) or (2) an increase/decrease of both rates (destructive; Fig. 3f).

## Discussion

CellRank 2 is a robust, modular, and scalable framework to infer and study single-cell trajectories and fate decisions. By separating the inference and analysis of transition matrices via *kernels* and *estimators*, respectively, CellRank 2 accommodates diverse data modalities and overcomes the limitations of single data-type approaches in a consistent and unified manner. Our tool successfully performed pseudotime-based analysis of human hematopoiesis and deciphered gene dynamics during human endoderm development using stemness estimates. Importantly, the modular and scalable design facilitated the rapid integration of each data modality and allowed CellRank 2 to analyze much larger datasets compared to previous approaches and implementations.

Developing an efficient optimal-transport-based kernel allowed us to integrate time series data, considering both inter and intra-time point information. With this formulation, we identified a putative progenitor population of medullary thymic epithelial cells missed by methods that ignore dynamics within time points. Recently, time course studies have been combined with genetic lineage tracing^57–60^ or spatial resolution^61–63^, and emerging computational methods^28, 29, 64–66^ use this information to map cells more faithfully across time. These enhanced inter-time point mappings can be used with our *RealTimeKernel* for further analysis, as demonstrated for lineage-traced C. elegans data in moslin^28^ and spatio-temporal mouse embryogenesis data in moscot^29^. These applications highlight the importance of our view-agnostic framework for analyzing increasingly large, complex, and multi-modal time course studies. Additionally, Mellon^67^, a recently proposed alternative approach for continuous analysis of time course data, could improve our mappings by incorporating their density estimates in the optimal transport problem.

Our kernel-estimator design proved particularly valuable when integrating metabolic labeling to estimate cell-specific mRNA transcription and degradation rates. We demonstrated the ability of metabolic labeling data to overcome the intrinsic limitations of splicing-based velocity inference by successfully identifying all lineages in gut organoid differentiation. Combining the inferred kinetic rates with CellRank 2 also makes it possible to study gene regulatory strategies underlying cellular state changes, as we showed for the goblet and enterocyte lineages. Parallel to our approach, Maizels *et al.* and Peng *et al.* developed velvet^68^ and storm^69^, respectively, to estimate cellular dynamics from metabolic labeling data. However, compared to our approach, velvet does not estimate transcription rates and assumes constant degradation rates across all cells. While storm relaxes this assumption, it does so only through post-processing steps. Additionally, storm relies on deterministic downstream analyses. In contrast, CellRank 2 estimates cell-specific transcription and degradation rates and offers probabilistic downstream analysis through flexible Markov-chain modeling.

Recent experimental advances combine single-cell metabolic labeling techniques with droplet-based assays^14, 70, 71^ or split-pool barcoding approaches^17, 68^ to label transcripts at atlas scale and demonstrate metabolic labeling for vivo systems^71, 72^ and in the context of spatially resolved assays^21^, underscoring the need for scalable analytical approaches as proposed in this study. We aim to expand our framework further by simultaneously inferring kinetic rates and ordering cells along differentiation trajectories.

We have introduced kernels that make use of different types of directional information of cellular state changes (Supplementary Fig. 1). If metabolic labels from pulse (chase) experiments for at least two (three) labeling durations are available, our proposed method to infer a metabolic-labeling informed vector field is ideally used. The *RealTimeKernel* is applicable for time series in which time points are closely spaced with respect to the underlying dynamical process. The *VelocityKernel* can be used with RNA velocity for systems that meet the assumptions of RNA velocity inference methods^73^. Finally, the *PseudotimeKernel* can enhance the understanding of cellular state changes if a unique initial state is identifiable and differentiation proceeds unidirectionally, and the *CyoTRACEKernel* can be used when the initial state is unknown. Different kernels can be combined with user-defined global weighting if multiple criteria are met, as we demonstrated for the *RealTimeKernel,* and we plan to introduce local kernel combinations in the future. Such an approach would involve kernel weights based on the relative position of cells within the phenotypic manifold allowing for context-dependent integration of multiple data sources.

Identifying driver genes is another aspect that can be extended in future work. Currently, we rank driver genes by correlating fate probabilities with gene expression. Although this approach has proven powerful, as shown in various applications, it is solely based on correlation. To unravel the causal mechanisms linking molecular properties and changes to fate decisions, perturbational data and causal inference^74^ can be combined with CellRank 2. This combination will ultimately enhance our understanding of underlying molecular drivers. Overall, we anticipate our framework to be crucial in understanding and conceptualizing fate choice as single-cell datasets grow in scale and diversity.

## People

We thank Nicolas Battich for helpful discussions about scEU-seq data, Rene Maehr and Macrina Lobo for insights into pharyngeal organ development, and T. Nawy for helping to write the manuscript. We would also like to thank Isaac Virshup, and numerous researchers who provided feedback to our implementation on GitHub.

## Funding

This work was supported by the BMBF-funded de.NBI Cloud within the German Network for Bioinformatics Infrastructure (de.NBI) (031A532B, 031A533A, 031A533B, 031A534A, 031A535A, 031A537A, 031A537B, 031A537C, 031A537D, 031A538A) and co-funded by the European Union (ERC, DeepCell - 101054957). M.L. acknowledges financial support from the Joachim Herz Foundation. F.J.T. acknowledges support by Wellcome Leap as part of the ΔTissue Program.

For all support coming via EU funding, the views and opinions expressed are those of the author(s) only and do not necessarily reflect those of the European Union or the European Research Council. Neither the European Union nor the granting authority can be held responsible for them.

## Author’s contributions

P.W. and M.L. contributed equally. P.W., M.L., and F.J.T. conceptualized the study. M.K. implemented CellRank 2 with contributions from P.W. and M.L.. P.W. designed and implemented the inference scheme for metabolic labeling data. P.W. performed analyses with input from M.L.. P.W., M.L., F.J.T, and D.P. wrote the manuscript. All authors read and approved the final manuscript.

## Competing interests

F.J.T. consults for Immunai Inc., Singularity Bio B.V., CytoReason Ltd, Cellarity, and Omniscope Ltd, and has ownership interest in Dermagnostix GmbH and Cellarity. D.P. is on the scientific advisory board of Insitro. The remaining authors declare no competing interests.

## Data availability

All data presented in this study is publicly available via the original publications.

## Code availability

CellRank 2 is released under the BSD-3-Clause license, with code available at https://github.com/theislab/cellrank. The inference of kinetic rates based on metabolic labeling data is implemented as part of the scVelo package: https://github.com/theislab/scvelo. Code to reproduce the results in the manuscript can be found at https://github.com/theislab/cellrank2_reproducibility.

## Supplementary Materials

Supplementary Figures: Supplementary Figures 1-11

Supplementary Notes: Supplementary Note 1

Supplementary Tables: Supplementary Table 1

## Supplementary Figures

**Suppl. Fig. 1.**
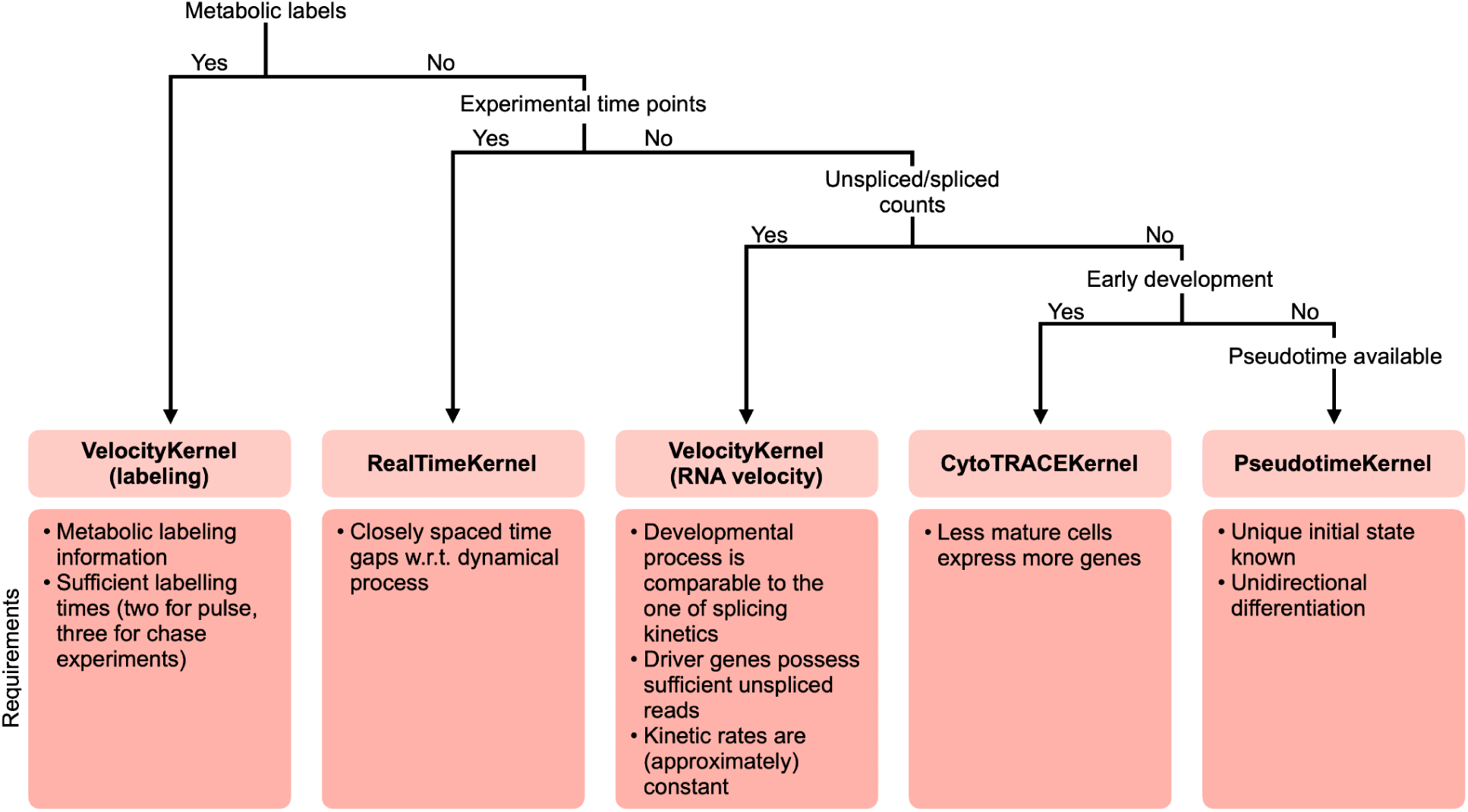
| Guiding kernel choice in CellRank 2. CellRank 2 implements various kernels which are suitable for different data modalities and experimental designs. The decision tree may be used as a guide to identify the most suitable kernel.

**Suppl. Fig. 2.**
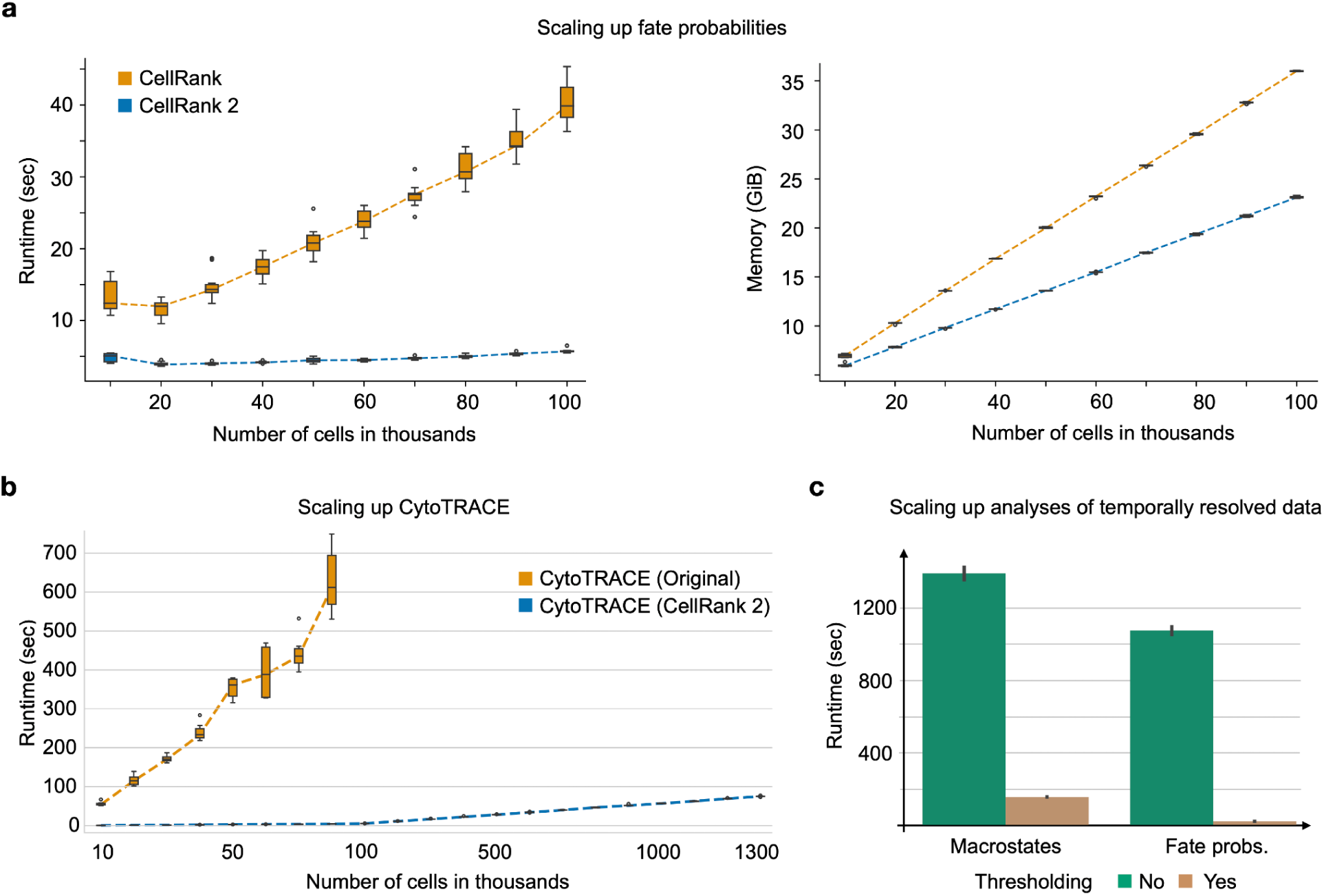
| CellRank 2 scales to large cell numbers. **a.** Runtime (left) and peak memory consumption (right) to compute fate probabilities with CellRank (orange) and CellRank 2 (blue). Both methods were run on subsets of a reprogramming dataset containing over 100,000 cells^75^. Box plots indicate the median (center line), interquartile range (hinges), and whiskers at 1.5x interquartile range (N=10 runs each). **b.** The CellRank 2 adaptation of CytoTRACE scales to a mouse organogenesis atlas of 1.3 million cells^8^, whereas CytoTRACE fails above 80,000 cells. Box plots indicate the median (center line), interquartile range (hinges), and 1.5x interquartile range (whiskers) (50,000 cells, original: N=6 runs; 60,000 cells, original: N=8 runs; 80,000 cells, original: N=9 runs; otherwise: N=10 runs). **c.** Runtime for calculating macrostates and fate probabilities using the *RealTimeKernel* with (brown) and without (green) thresholded transition matrix. The mean execution time of 10 runs and the corresponding 95% confidence interval is shown.

**Suppl. Fig. 3.**
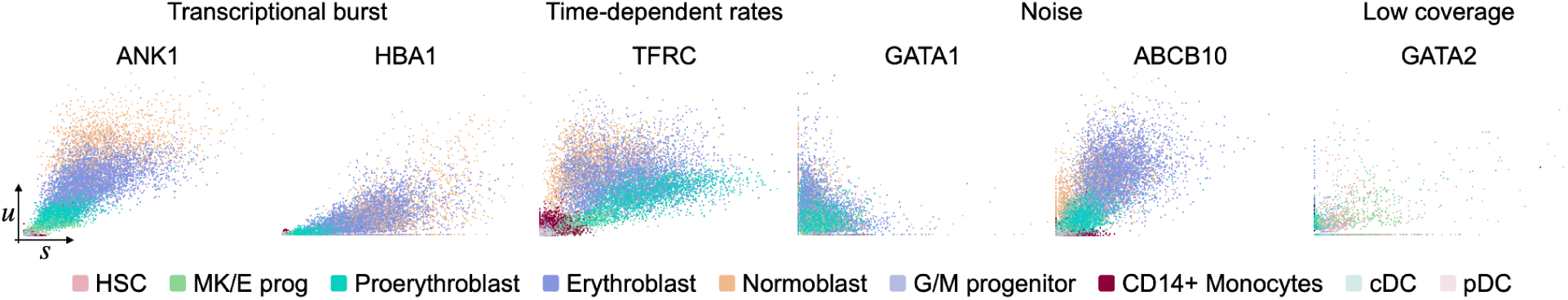
| RNA velocity model assumptions violate transcriptional kinetics in adult hematopoiesis. Exemplary phase portraits of genes violating assumptions of the inference algorithm in scVelo’s *dynamical model*^12^ on the adult hematopoiesis dataset of Fig. 1. Transcriptional bursts (ANK1, HBA1) and time-dependent kinetic rates (TFRC) violate scVelo’s constant-rate assumption. Very high noise levels (GATA1, ABCB10) and low coverage in either spliced or unspliced transcripts (GATA2) render the inference problem ambiguous (Supplementary Note 1). Each dot is a cell colored by cell type (HSC: hematopoietic stem cell, MK/E prog: megakaryocyte/erythrocyte progenitors, G/M progenitor: Granulocyte/Myeloid progenitor, pDC: plasmacytoid dendritic cell, cDC2: classical dendritic cells) according to original cluster annotations^7^; the x-axis (y-axis) shows the abundance of spliced (unspliced) counts.

**Suppl. Fig. 4.**
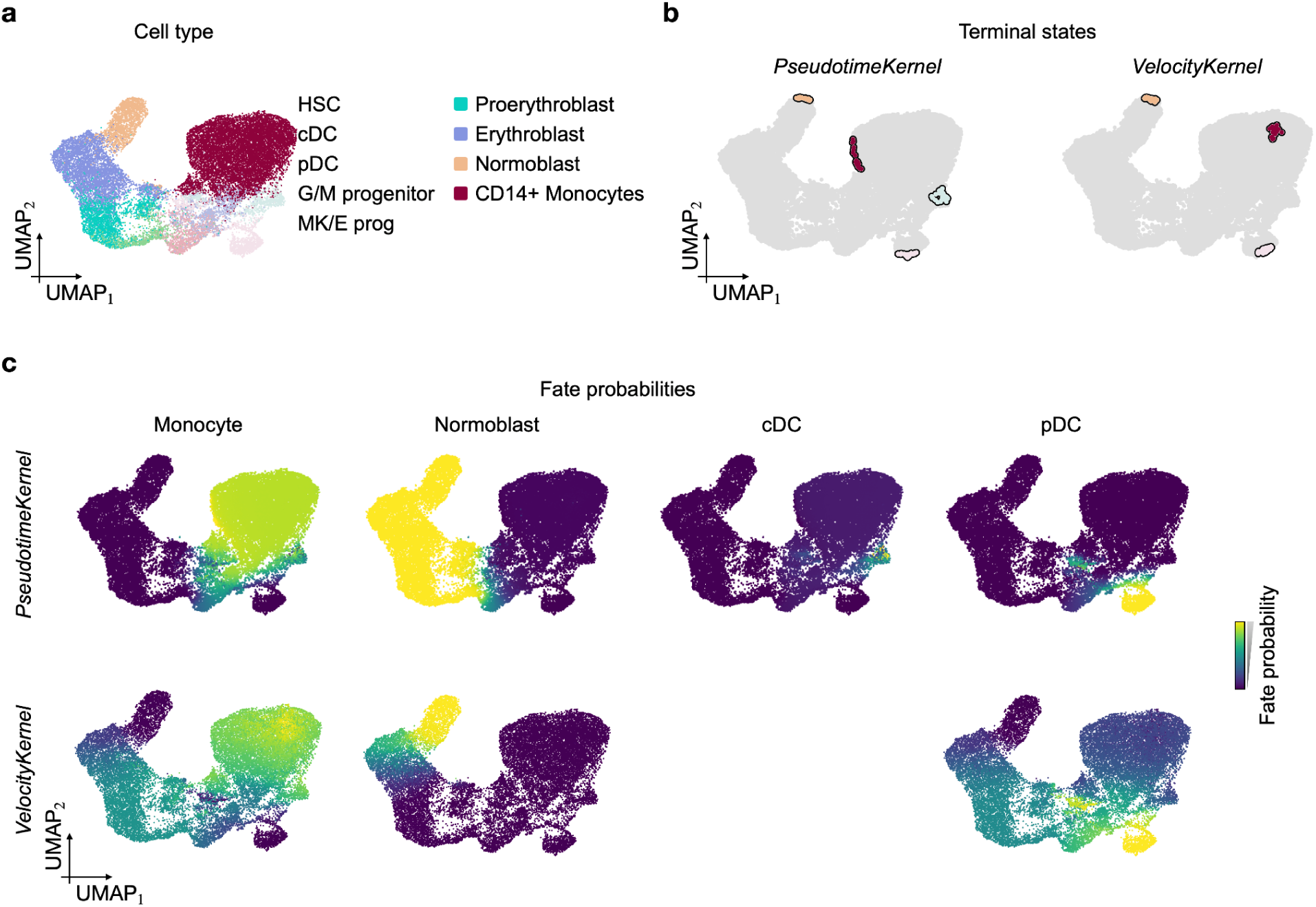
| Fate mapping in hematopoiesis with the *PseudotimeKernel* and *VelocityKernel*. **a.** UMAP embedding of entire hematopoiesis dataset^31^. Cell types are colored according to the original publication. The data subset of interest is highlighted in black. **b.** Terminal states identified by the *PseudotimeKernel* (top) and *VelocityKernel* with RNA velocity (bottom) inferred using scVelo’s *dynamical model*^12^. **c.** Fate probabilities towards each identified terminal state based on the *PseudotimeKernel* (top) and *VelocityKernel* (bottom). The *VelocityKernel* does not identify the cDC terminal state.

**Suppl. Fig. 5.**
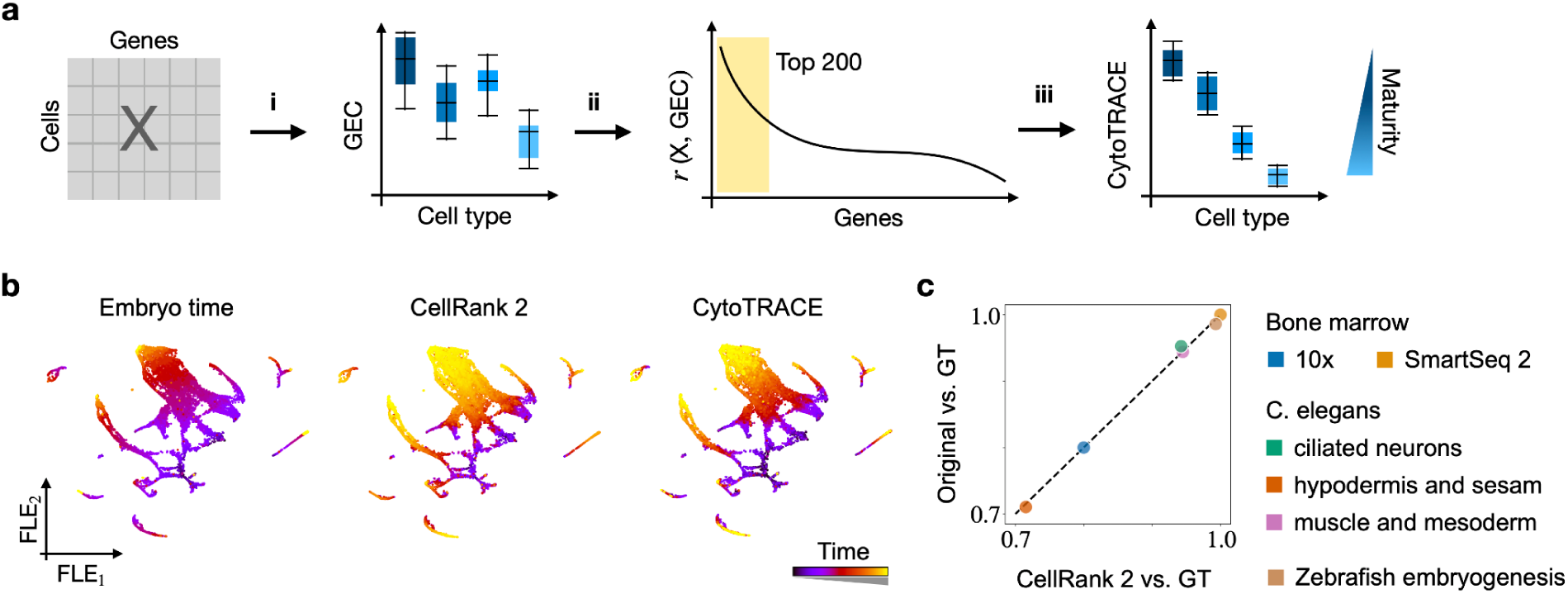
| Developing the *CytoTRACEKernel*. a. Similar to the CytoTRACE publication^25^, we compute the CytoTRACE score by (i) calculating the number of genes expressed per cell (GEC), (ii) computing each gene’s Pearson correlation with GEC, (iii) mean-aggregating imputed expression of the top 200 correlated genes (Methods). b. Force-directed layout embedding (FLE) of 22,370 Caenorhabditis (C.) elegans muscle and mesoderm cells undergoing embryogenesis, colored by estimated embryo time^76^ (left; 130-830 min), CytoTRACE pseudotime computed using CellRank 2 (middle) and the original implementation (right). c. Quantitative comparison of the two implementations of the CytoTRACE pseudotime on bone marrow^77^ (using 10x and SmartSeq2), C. elegans embryogenesis^76^ (subsetted to ciliated neurons, hypodermis and seam, and muscle and mesoderm), and zebrafish embryogenesis^5^. The x-axis (y-axis) displays Spearman’s rank correlation between CellRank 2-CytoTRACE (original CytoTRACE) and ground truth (GT) time labels. Ground truth labels were derived from either embryo time or stages as in b (C. elegans and zebrafish embryogenesis), or from manually assigned maturation labels from the original CytoTRACE study^25^ (bone marrow).

**Suppl. Fig. 6.**
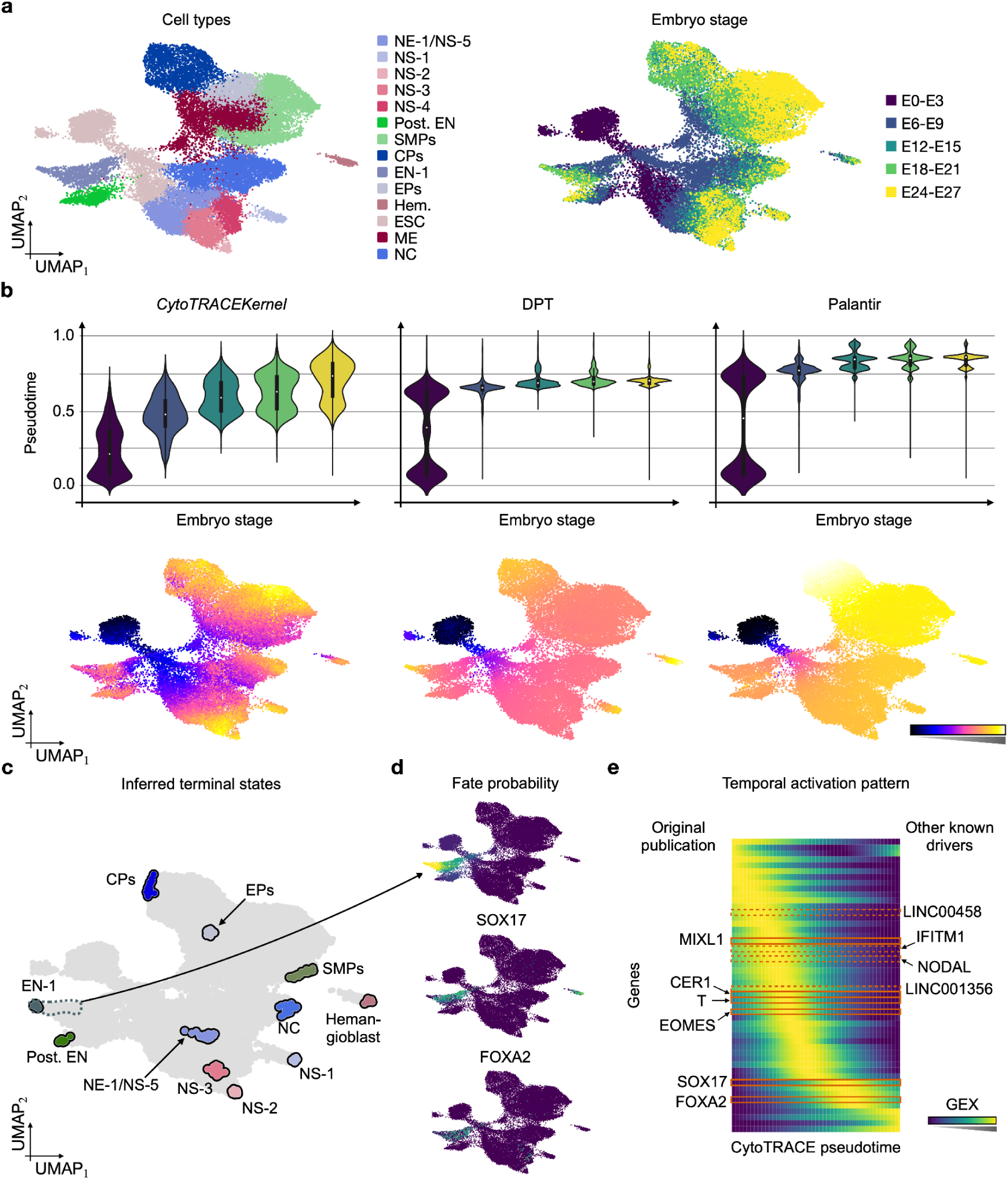
| The *CytoTRACEKernel* recovers temporal gene activation. **a**. UMAP embedding of 31,029 embryoid-body cells^35^, colored by original cluster annotation (left) and embryo stage (right). **b.** Distribution of pseudotimes from the *CytoTRACEKernel* (left), DPT (center) and Palantir (right), stratified by embryo stage (top row) and colored according to **a**. Box plots indicate the median (center line), interquartile range (hinges), and 1.5x interquartile range (whiskers) (E0-E3: N=4574 cells, E6-E9: N=7368 cells, E12-E15: N=6241 cells, E18-E21: N=6543 cells, E24-E27: N=6302 cells). UMAP embeddings (bottom) are colored by pseudotime. c. Terminal states inferred using the CytoTRACEKernel. d. UMAP embedding colored by fate probabilities (top) and recovered driver gene expression (bottom). e. Smoothed gene expression along the CytoTRACE pseudotime for the automatically identified top 50 driver genes, sorted according to their pseudotime peak. Genes identified in the original publication (left) and known drivers (right)35–38 additionally recovered with CellRank 2 are indicated.

**Suppl. Fig. 7.**
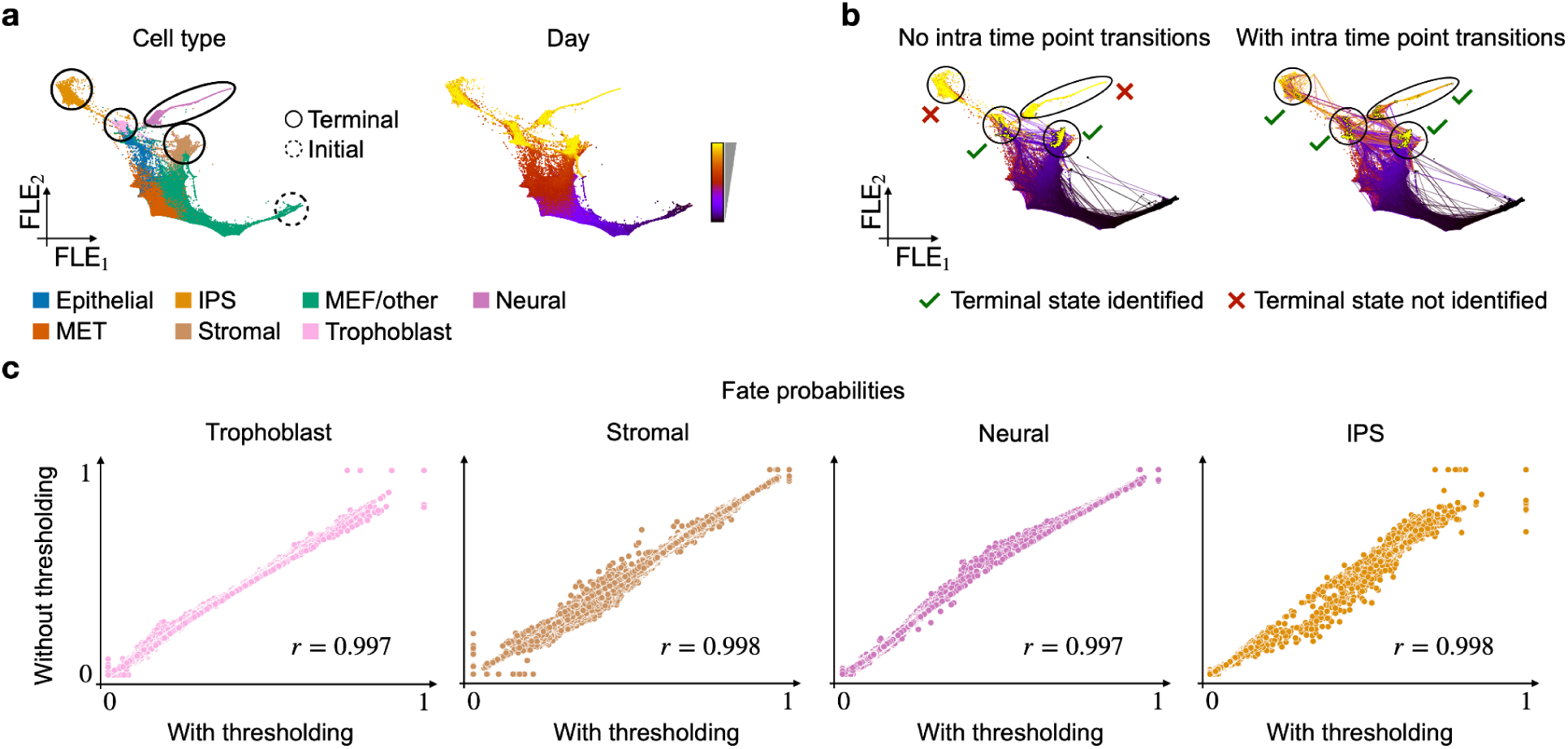
| Developing the *RealTimeKernel*. **a**. Force-directed layout embedding (FLE) of 165,892 mouse embryonic fibroblasts (MEFs) reprogramming towards various endpoints during an 18-day time course^23^, colored according to modified original annotations (IPS: induced pluripotent stem; left) or sequencing time points (right). Dotted (solid) circles indicate known initial (terminal) states. **b.** FLE showing simulated random walks from day 0 cells without (left; corresponds to WOT^23^) and with (right; corresponds to the *RealTimeKernel*) intra-time point transitions. Black (yellow) dots denote a random walk’s start (end). Green ticks (red crosses) indicate known terminal states which are (are not) explored by random walks. **c.** Evaluation of the effect of thresholding the transition matrix in the *RealTimeKernel*. Each dot corresponds to one cell’s fate probability towards one of four terminal states, computed with thresholding (x-axis) and without thresholding (y-axis). The color coding is in agreement with **a**.

**Suppl. Fig. 8.**
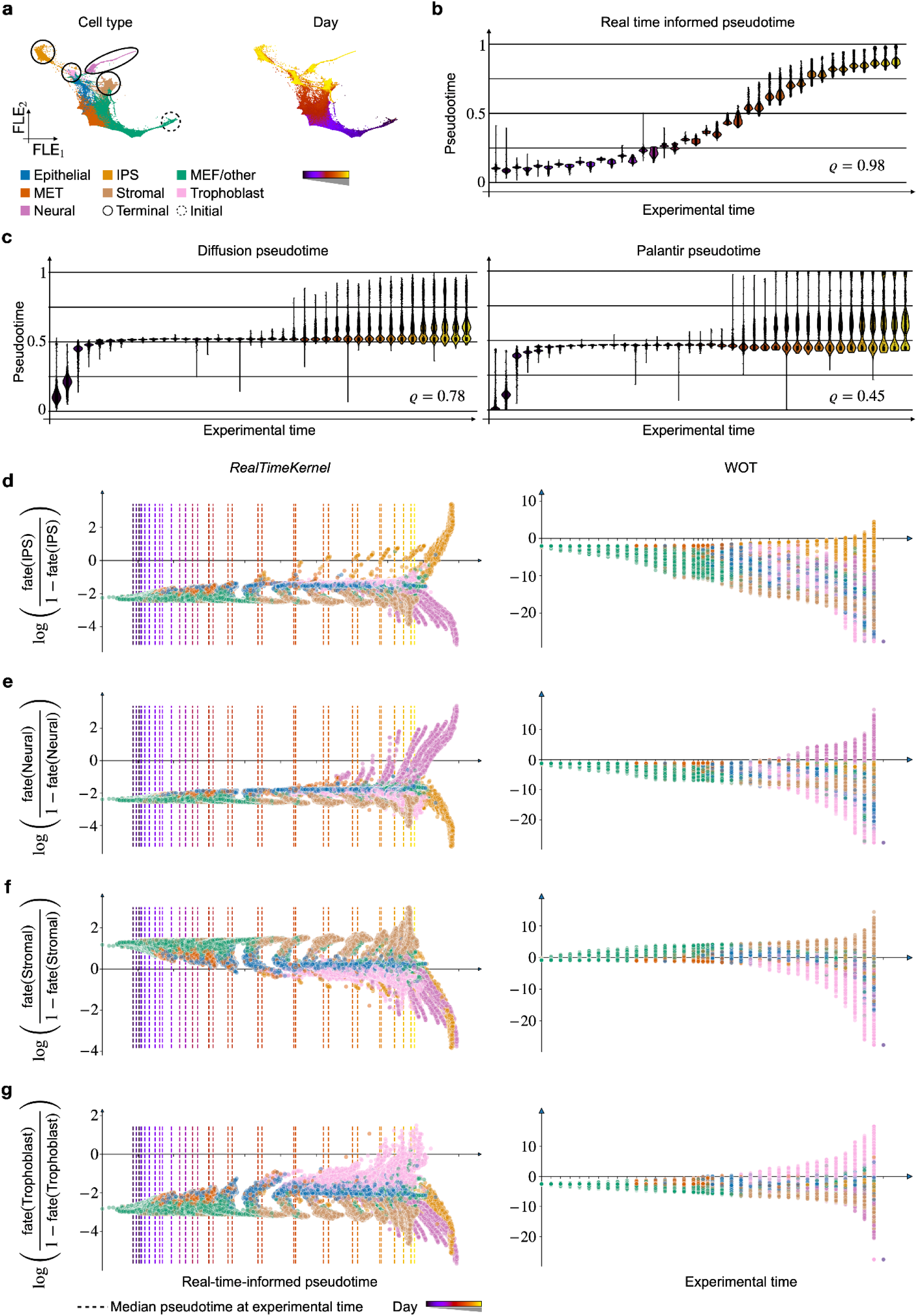
| The *RealTimeKernel* studies fate decision on a continuous domain. **a**. Force-directed layout embedding (FLE) of 165,892 mouse embryonic fibroblasts (MEFs) reprogramming towards various endpoints during an 18-day time course^23^, colored according to modified original annotations (IPS: induced pluripotent stem; left) or sequencing time points (right). Dotted (solid) circles indicate known initial (terminal) states. **b,c.** Violin plots showing pseudotime distribution for each of 36 experimental time points, using CellRank 2’s real-time-informed pseudotime (**b.**), DPT^2^ (**c.**, left), or Palantir ^7^ (**c.**, right) (Methods). Box plots indicate the median (center line), interquartile range (hinges), and whiskers at 1.5x interquartile range. **c.** DPT^2^ (center), or Palantir ^7^ (right). The number of cells per experimental time point is specified in Supplementary Table 1. **d-g.** Cell-specific fate change over pseudotime (*RealTimeKernel*, left) or experimental time (WOT, right) for the IPS (**d.**), neural (**e.**), stromal (**f.**), and trophoblast lineage (**g.**). For the *RealTimeKernel*, dashed vertical lines denote the mean pseudotime over all cells from a given experimental time point, recapitulating the correct ordering from **b**.

**Suppl. Fig. 9.**
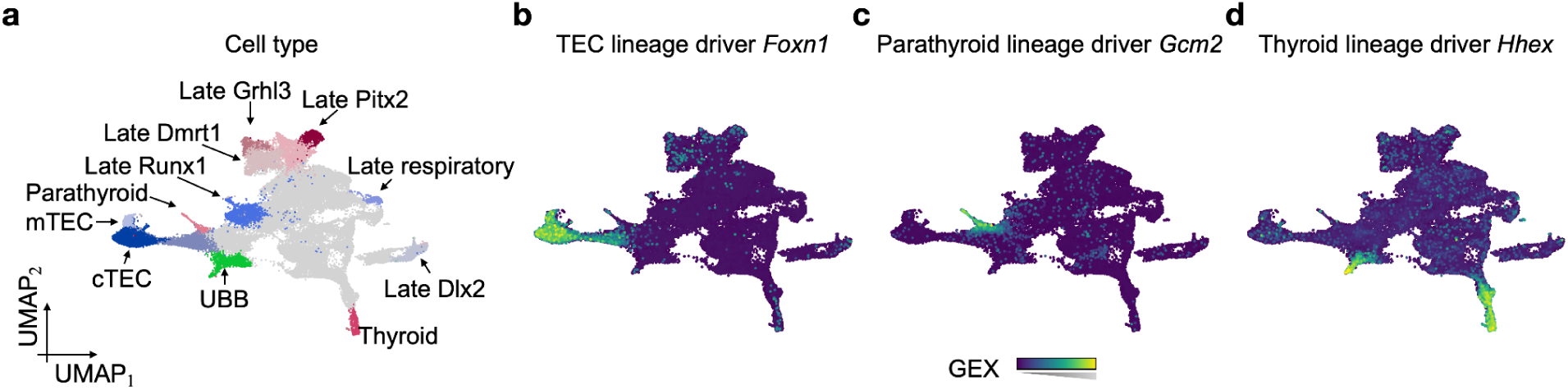
| Driver genes of pharyngeal endoderm development. **a.** UMAP embedding of pharyngeal endoderm development dataset. Cells are colored according to the original study. **b-d.** UMAP embedding colored by CellRank 2-identified lineage drivers *Foxn1* (TEC, **b.**), *Gcm2* (parathyroid; **c.**), *Hhex* (thyroid; **d.**).

**Suppl. Fig. 10.**
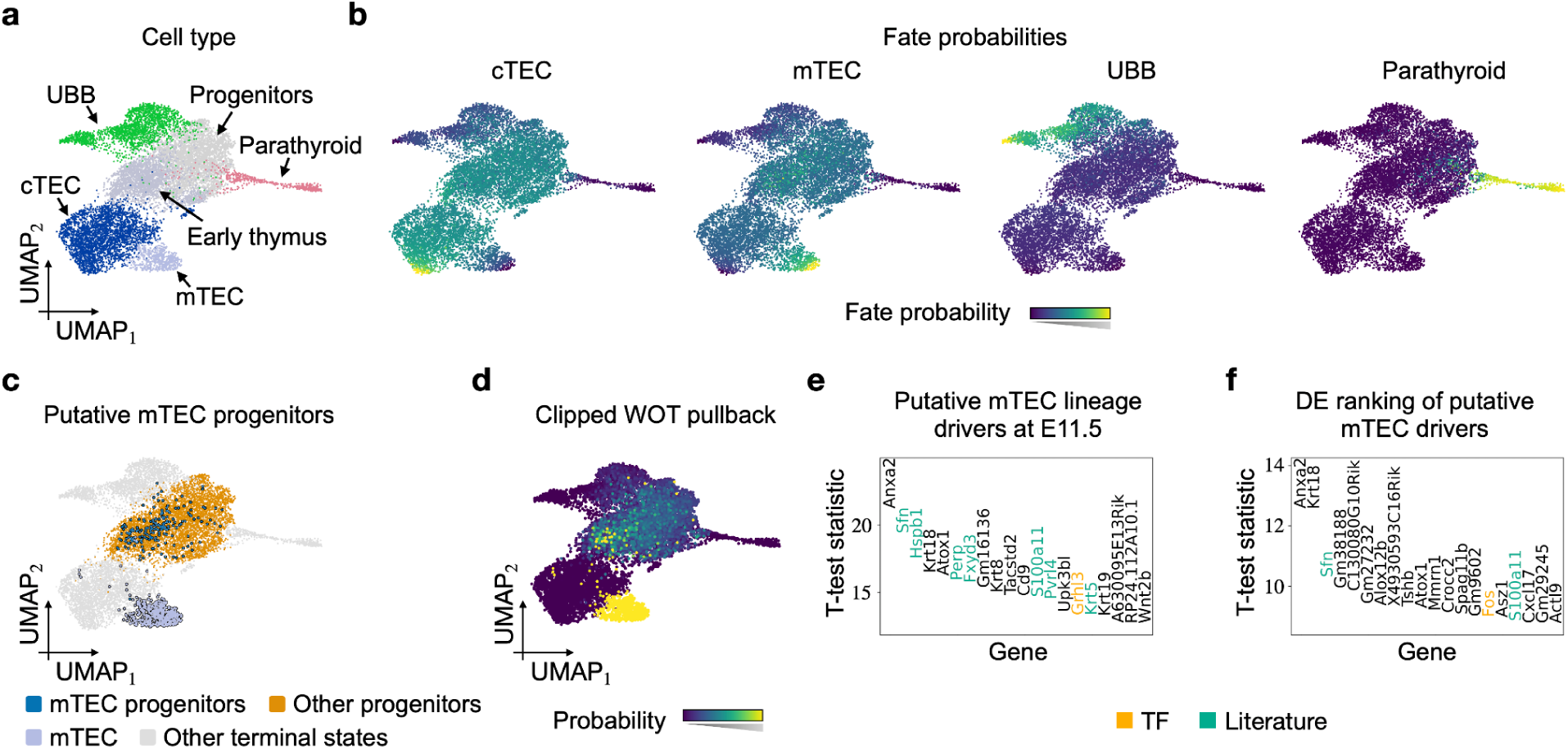
| Putative mTEC progenitors. **a.** UMAP embedding of subsetted pharyngeal endoderm dataset. Cells are colored according to the original study. **b.** The fate probabilities towards each terminal state are estimated using the *RealTimeKernel*. **c.** A cluster of putative mTEC progenitors is identified by cells with fate probabilities towards mTEC larger than 0.5. **d.** UMAP embedding colored by WOT’s pullback at E10.5 clipped to the 99 percentile. **e.** Gene ranking of potential mTEC drivers based on WOT’s pullback distribution at E11.5, compared to the gene ranking based on the pullback at E10.5, as shown in Fig. 2f. **f.** Gene ranking when performing classical differential expression analysis on clusters “mTEC progenitors” and “Other progenitors” as shown in **c.**

**Suppl. Fig. 11.**
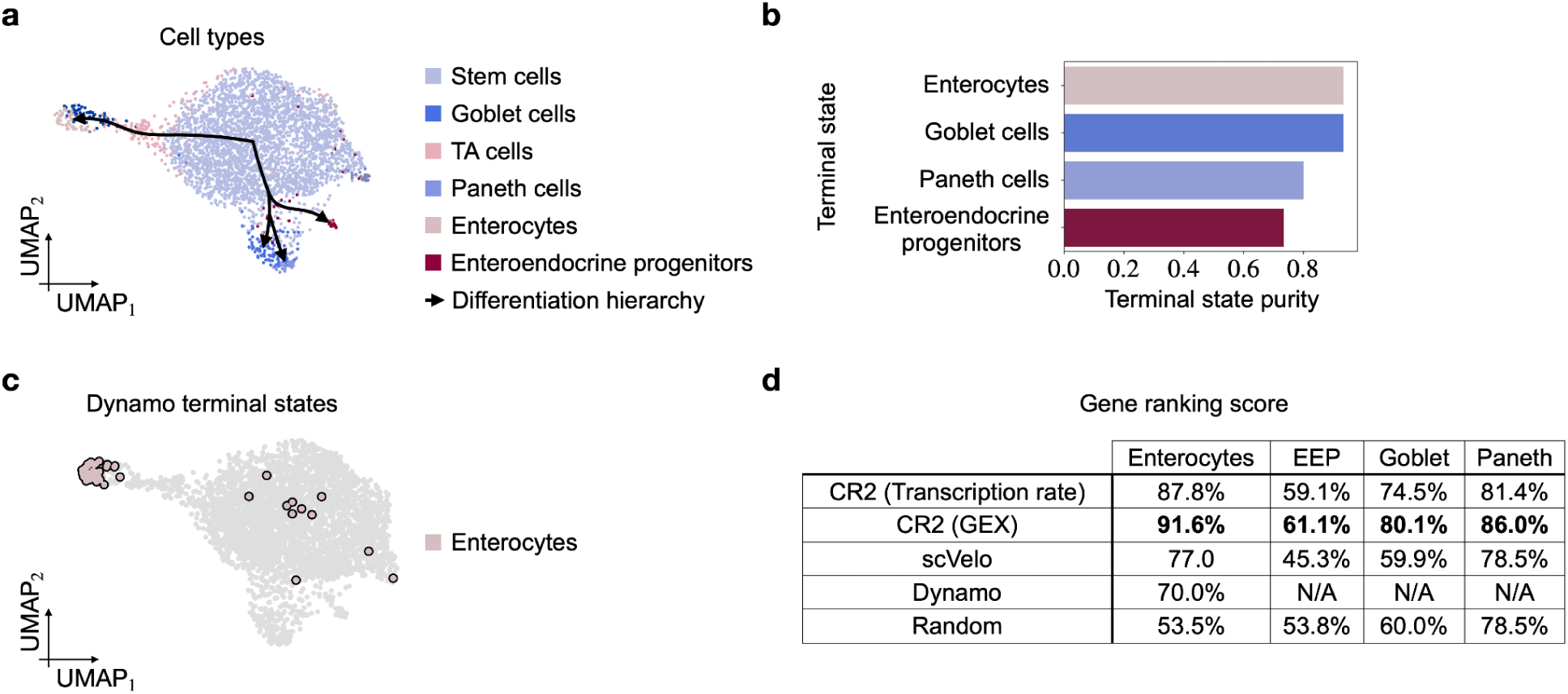
| Terminal state analysis for metabolically labeled intestinal organoids. **a**. UMAP embedding of intestinal cells. Cells are colored according to their cell type (TA cells: transit-amplifying cells) according to the original publication^15^. Black arrows indicate the known differentiation hierarchy. **b.** Terminal state purity identified using our metabolic labeling-based vector field. **c.** UMAP embedding highlighting terminal states identified by dynamo^24^. **d.** The ranking score metric for each method and terminal state. The metric quantifies the degree of optimality of a gene ranking compared to an optimal ranking (CR2: CellRank 2). Dynamo only identified the trajectory leading up to enterocytes.

## Methods

### CellRank 2: A unified framework to probabilistically model cellular state changes

To overcome the inherent limitations of RNA velocity and unify trajectory inference across different data views, we developed CellRank 2. Our framework describes cellular dynamics probabilistically, as proposed in our earlier work^22^. Specifically, we introduced the first probabilistic modeling framework that automatically determines the direction of cellular state changes to extend trajectory inference beyond normal development. Generalizing this paradigm to different biological priors and guaranteeing applicability in many scenarios required us to rethink the CellRank structure entirely. To this end, we base our new version on three key principles:

1. **Robustness:** Fate restriction is a gradual, noisy process requiring probabilistic treatment. Therefore, we use Markov chains to describe stochastic fate transitions, with each state of the Markov chain representing one cell.
2. **Modularity:** Quantifying transition probabilities between cells is independent of analyzing them. Thus, we modularized the CellRank framework into *kernels* to compute transition probabilities, and *estimators* to analyze transition probabilities. This structure guarantees flexibility in applications and is easily extensible.
3. **Scalability:** We assume each cell can transition into a small set of possible descendant states. Consequently, transition matrices are sparse, and computations scale to vast cell numbers (Supplementary Fig. 2).

### Innovations in CellRank 2

Our design principles allowed us to improve our original work in three major aspects:

1. We design a modular interface allowing us to decouple the construction of a Markov chain from the process of formulating a hypothesis based on the Markov chain.
2. We introduce the *PseudotimeKernel*, *CytoTRACEKernel*, and *RealTimeKernel*, as well as a method to infer kinetic rates from metabolic labeling data, to render CellRank 2 applicable beyond RNA velocity by using a pseudotime, a measure of developmental potential, time-course data, and metabolic labeling information, respectively.
3. We make our framework faster by accelerating our main estimator by one order of magnitude, easier to use by refactoring our codebase, and more interpretable by visualizing kernel dynamics via random walks.

### Key outputs of CellRank 2

Although inputs to CellRank 2 are kernel-dependent (Supplementary Fig. 1), outputs are consistent across all kernels:

● Initial, intermediate, and terminal states of cellular trajectories.
● Fate probabilities, quantifying how likely each cell is to reach each terminal (or intermediate) state.
● Gene expression trends specific to each identified trajectory.
● Putative driver genes of fate decisions through correlating gene expression with fate probability.
● Dedicated visualization tools for all key outputs, *e.g.*, circular embeddings for fate probabilities, heatmaps for cascades of trajectory-specific gene expression, and line plots for gene trends along different trajectories.

### A conceptual overview of kernels in CellRank 2

#### Decoupling inference of transition probability from their analysis

The typical CellRank 2workflow consists of two steps: (1) estimating cell-cell transition probabilities and (2) deriving biological insights based on these estimates. Previously, we tied these two steps together^22^ but realized that decoupling them yields a much more powerful and flexible modeling framework. Treating each step separately is possible since analyzing transition matrices is independent of their construction. For example, estimating transition probabilities based on RNA velocity or a pseudotime does not change how initial and terminal states are inferred or fate probabilities estimated. Consequently, modularizing our problem-specific framework generalizes the corresponding analysis tools to other data modalities. The two steps of our inference workflow are conceptualized by *kernels* and *estimators*, respectively.

***Kernels*** estimate transition matrices *T* ∈ ℝ^*n_c_* × *n_c_*^ at a cellular resolution with *n_c_* denoting the number of cells. Each row *T_j_*,: represents the transition probabilities of cell towards putative descendants. With CellRank 2, we provide means to quantify fate probabilities based on RNA velocity (*VelocityKernel*), pseudotime (*PseudotimeKernel*), a developmental potential (*CytoTRACEKernel*), experimental time points (*RealTimeKernel*), and metabolic labeling (metabolic-labeling-based vector field with the *VelocityKernel*).

### Kernel combination

Different data modalities may capture different aspects of biological processes. To take advantage of multiple data modalities, *kernels* can be combined to quantify the likely state change in a single, aggregated transition matrix. Consider two kernels *k*^(1)^ and *k*^(2)^ with corresponding transition matrices *T*^(1)^ and *k*^(2)^, respectively. CellRank 2 allows combining the two kernels into a joint kernel *k* defined as *k* = α*k*^(1)^ + (1 − α)*k*^(2)^ with a weight parameter α ∈ [0,1]. The corresponding normalized transition matrix is computed automatically and is, thus, given by

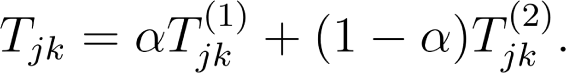

#### Visualizing kernel dynamics: Random walks and projections

Although inferred transition probabilities are predominantly used for more in-depth data analyses based on *estimators*, we also provide means to visualize cellular dynamics directly based on the kernel output. These visualizations are intended to provide a preliminary understanding of the underlying dynamics and serve as a starting point for further analyses. Here, we enable studying the evolution of cellular state change either based on random walks in the high-dimensional gene expression space or a projection of the high-dimensional vector field onto a low-dimensional latent space representation of the data.

Transition matrices induce random walks modeling the evolution of individual cells. Given a cell *j*, we successively sample its future state under the given transition matrix. Starting cells for random walks can be sampled either at random or from a user-defined early cell cluster. We terminate random walks when a pre-defined maximum number of steps has been performed or when a predefined set of terminal cells has been reached. By studying multiple random walks, the expected dynamics are revealed. Random walks, including their start and final cells, can then be visualized in a low-dimensional representation of the data. Within our framework, random walks are computed efficiently via a parallel implementation.

Previously, the most popular approach for visualizing RNA velocity has been the projection of the high-dimensional vector field onto a low-dimensional latent space representation^13^. With CellRank 2, we generalize this concept to any *kernel* based on a k-nearest neighbor graph, *i.e.*, the *PseudotimeKernel*, *CytoTRACEKernel*, and *VelocityKernel*. The projection for a given cell is calculated as follows: Consider a transition matrix *T*, cell *j* with neighborhood *N*(*j*) and *k_j_* neighbors, and latent representation *z_j_*. The projected velocity *v_j_* is then given by

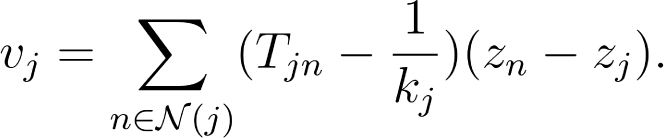

While we provide the option to visualize the projected velocity stream in low dimensions for specific kernels, we caution against the analysis thereof. Previous work^78, 79^ highlighted how the projected velocity stream is sensitive to many parameters, including the gene set, the embedding technique, and more (Supplementary Note 1). Instead, we encourage visualizing cellular dynamics through random walks, sampled independently of the embedding, or through initial and terminal states, fate probabilities, and other quantities inferred in high dimensions through our estimator modules.

### A conceptual overview of estimators in CellRank 2

Based on transition matrices provided by *kernels*, we enable data-driven knowledge discovery. To this end, *estimators* first identify initial, intermediate, and terminal states using the pre-computed transition matrices. States are identified using concepts and results from the rich theory of Markov chains. Following, we enable visualizing trajectory-specific gene expression trends and cascades of gene activation^22^, clustering expression trends^22^, or arranging cells in a circular embedding^22, 80, 81^ to summarize fate probabilities. We provide the necessary tools for each step of the downstream analysis as part of CellRank 2.

#### The GPCCA estimator

As in our previous work, we compute macrostates and classify them as initial, intermediate, and terminal by coarse-graining the cell-cell transition matrix. This approach is based on *Generalized Perron Cluster Cluster Analysis*^26, 27, 82^ *(GPCCA)*, a method initially developed to study conformational protein dynamics.

### Performance improvements of CellRank 2

#### Faster computation of fate probabilities

After estimating cell-cell transition probabilities through a *kernel* and identifying terminal states through an *estimator*, we assess cellular fate towards these terminal states. For each cell, we quantify its fate probability, *i.e.*, how likely it is to differentiate into one of the terminal states. Given our Markov-chain-based framework, fate probabilities can be computed in closed form using absorption probabilities. However, calculating absorption probabilities directly scales cubically in the number of cells. To overcome this computational burden, in our previous work, we reformulated the underlying problem as a set of linear systems. These linear systems are then solved in parallel using a sparsity-optimized iterative algorithm^83^. This reformulation scales near-linearly^22^.

Even though our previously proposed reformulation for computing absorption probabilities achieved a significant increase in performance compared to a naive implementation, we still encountered increased runtimes when analyzing larger datasets (Supplementary Fig. 2a). To reduce the runtime further, we devised an alternative but equivalent approach: Given a terminal state, we previously identified *n_f_* representative cells, computed absorption probabilities towards them, and aggregated them across the *n_f_* representative cells to assign a single, lineage-specific probability. In CellRank 2, we first combine the *n_f_* representative cells into a single pseudo-state and compute absorption probabilities towards it instead. While the corresponding results are mathematically equivalent, ignoring parallelization, this new approach is *n_f_* times faster. Therefore, with *n_f_* = 30 by default, our improved implementation results in a 30-fold speedup.

#### Extensibility of CellRank 2

While we already provide multiple *kernels* tailored to different data modalities, current and future technologies provide additional sources of information. Concrete examples include spatially resolved time course studies^61–63^ and genetic lineage tracing data^57–60^, previously already integrated in the CellRank 2 ecosytem^28, 29^. Our modular interface makes CellRank 2 easily extensible towards (1) alternative single-cell data modalities by including new kernels, and (2) alternative trajectory descriptions generating different hypotheses through new estimators.

### The PseudotimeKernel: Incorporating prior knowledge on differentiation

Aligning cells along a continuous pseudotime mimicking the underlying differentiation process has been studied in many use cases. In particular, a pseudotime can be computed for systems where a single, known initial cellular state develops unidirectionally into a set of unknown terminal states. Based on the assigned pseudotime values, we quantify transition probabilities between cells using the *PseudotimeKernel*.

Given a similarity-based nearest neighbor graph with a corresponding adjacency matrix *C̃*, the *PseudotimeKernel* biases graph edges toward increasing pseudotime: Consider a reference cell *j*, one of its neighbors, *k* the corresponding edge weight Δ*C̃_jk_*, and the difference between their pseudotimes Δ*t_jk_*. To favor cellular transitions toward increasing pseudotime, the *PseudotimeKernel* downweighs graph edges pointing into the reference cell’s pseudotemporal past while leaving the remaining edges unchanged. Edge weights are updated according to

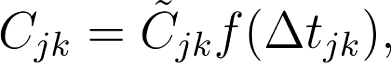

with a function *f* implementing the thresholding scheme. In CellRank 2, we implement soft and hard thresholding. The soft scheme continuously downweighs edge weights according to

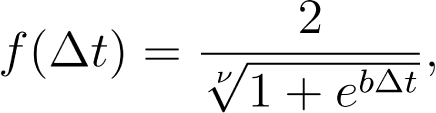

if Δ*t* < 0 and 1 otherwise. By default, the parameters *b* and *v* are set to 10 and 0.5, respectively. This concept is similar to the scheme proposed by the trajectory inference method VIA^30^. In contrast to soft thresholds, hard thresholding follows a stricter policy inspired by Palantir^7^ by discarding most edges that point into the pseudotemporal past.

### The CytoTRACEKernel: Inferring directionality from developmental potential

CytoTRACE assigns each cell in a given dataset a developmental potential^25^. Score values range from 0 to 1, with 0 and 1 identifying mature and naive cells, respectively. Inverting the score, thus, defines a pseudotime for developmental datasets. In CellRank 2, the *CytoTRACKernel* computes the CytoTRACE score and constructs the corresponding pseudotime to calculate a transition matrix as described for the *PseudotimeKernel*.

### Adaptation of the CytoTRACE score

When calculating the CytoTRACE score on larger datasets, we found the score construction either intractable due to long runtimes (40,000 to 80,000 cells) or failed to compute the score at all (more than 80,000 cells) (Supplementary Fig. 2b). Thus, to ensure computational efficiency to reconstruct the CytoTRACE score for larger datasets, we sought an alternative, computationally efficient and numerically highly correlated approach.

Conceptually, CytoTRACE proposes that the number of expressed genes decreases with cellular maturity. This assumption is biologically motivated by less developed cells regulating their chromatin less tightly^25^. The computation of the CytoTRACE score with CellRank 2 is composed of three main steps (Supplementary Fig. 5a). Consider the gene expression matrix and the smoothed gene expression matrix *X*^(smoothed)^ found by nearest-neighbor smoothing as implemented in scVelo^12^ or MAGIC^84^. For each cell *j*, we compute the number of genes it expresses (GEC ϵ ℕ^*n_c_*^), i.e.,

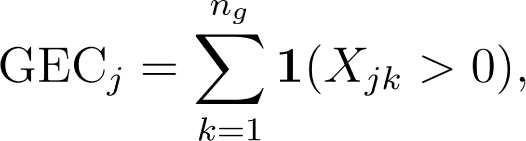

with indicator function 1(·). The indicator function equates to one if its argument holds true and zero otherwise. Next, for each gene, we compute its Pearson correlation with GEC, select the top *L* genes (default: 200), and subset *X*^smoothed)^ to the identified *L* genes. Finally, we mean-aggregate each cell’s gene expression

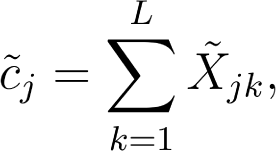

with *X̃* ϵ^*n_c_*×*L*^ denoting the subsetted, smoothed gene expression matrix *X*^(smoothed)^. The CytoTRACE score *c* is then given by scaling *c̃* to the unit interval

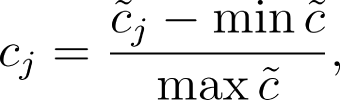

and the corresponding pseudotime *p*_cyt_ by inverting *c, i.e.*,

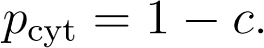

#### Comparison of the CytoTRACE score construction

Considering the nearest-neighbor-smoothed gene expression matrix, instead of an alternative, computationally more costly imputation scheme, is the main difference between our adaptation and the original CytoTRACE proposal. To impute gene expression, the original implementation solves a non-negative least squares regression problem and simulates a diffusion process^25^.

To confirm that our adapted scheme yields numerically similar results, we compared the CytoTRACE scores of the original and our approach to ground truth time or stage labels on six datasets previously used to validate CytoTRACE^25^ (Supplementary Fig. 5b,c). The considered datasets are bone marrow^77^ (using 10x and SmartSeq2), C. elegans embryogenesis^76^ (subsetted to ciliated neurons, hypodermis and seam, or muscle and mesoderm), and zebrafish embryogenesis^5^. For each dataset, the original CytoTRACE study derived ground truth time labels using either embryo time, stages (C. elegans and zebrafish embryogenesis), or a manual assignment (bone marrow). The concordance of each approach with ground truth was confirmed by calculating the Spearman rank correlation between the CytoTRACE score and ground truth time or stage labels.

### The *RealTimeKernel*: Resolving non-equilibria systems through time-series data

Commonly used single-cell sequencing protocols are destructive by design and offer, thus, only a discrete temporal resolution. Recent advances allow reconstructing transcriptomic changes across experimental time points using optimal transport (OT)^23^. However, these approaches focus only on inter-time-point information. Conversely, the *RealTimeKernel* incorporates both inter- and intra-time-point transitions to draw a more complete picture of cellular dynamics.

To quantify inter-time-point transitions, the *RealTimeKernel* relies on Waddington Optimal Transport^23^ (WOT). For each tuple of consecutive time points *t_j_* and *t_j+1_*, WOT identifies a transport map π_*t_j_*, *t_j_*+ 1_, assigning each cell at time *t_j_* its like future state at time *t_j+1_*. In addition, to study transcriptomic change within a single time point *t_j_*, we rely on transcriptomic similarity. We combine WOT-based inter-time point transport maps π*t_j_, t_j+1_* with similarity-based intra-time point transition matrices *T̃_t_j__,t_j_* in a global transition matrix which contains cells from all time points. In the global transition matrix, we place WOT-computed transport maps on the first off-diagonal, modeling transitions between subsequent time points, and similarity-based transition matrices on the diagonal, modeling transitions within each time point. We normalize each row to sum to one, giving rise to a Markov chain description of the system.

#### Thresholding transport maps for scalability

In the single-cell domain, most OT-based approaches, including WOT, rely on entropic regularization^41^ to speed up the computation of transport maps. However, entropic regularization leads to dense transport maps π_*t_j_*, *t_j_*+ 1_, rendering downstream computations based on the *RealTimeKernel* extremely expensive for larger datasets (Supplementary Fig. 7c), but most of the entries found in π_*t_j_*, *t_j_*+ 1_ are extremely small as a consequence of entropic regularization. As a result, these entries contribute only marginally to the observed dynamics (as we show below).

To ensure fast *RealTimeKernel*-based computations, we devised an adaptive thresholding scheme resulting in sparse transition matrices. Transition probabilities falling below a certain threshold are set to zero, all others are kept unchanged. Per default, we identify the smallest threshold that does not remove all transitions for any cell, *i.e.*,

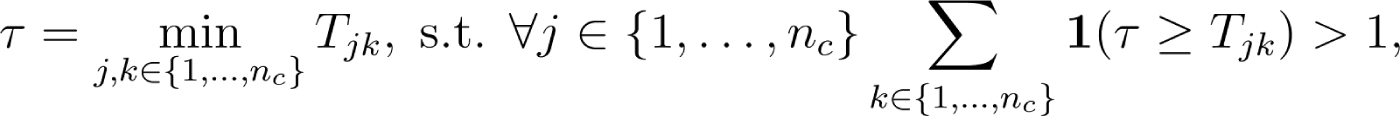

with indicator function 1(·). Alternatively, the same heuristic can be applied for each time point independently, or a user-defined threshold may be used. Following thresholding, we re-normalize the transition matrix such that rows sum to one again.

To verify that thresholding the transition matrix does not alter biological findings, we compared fate probabilities derived from the original and the thresholded transition matrix on a dataset of mouse embryonic fibroblast reprogramming^23^. For each terminal state, we computed the Pearson correlation between fate probabilities estimated by each approach.

### Estimating cellular fate from time-resolved scRNA-seq data

Traditional single-cell sequencing protocols include cell lysis and are, thus, destructive by nature. Consequently, the transcriptome can only be measured once, resulting in snapshot data. Recently, metabolic labeling approaches have been extended to single-cell resolution, providing an opportunity to overcome this challenge by measuring newly synthesized mRNA in a given time window^55^. To label transcripts, current protocols rely on the nucleoside analaogues 4-thiouridine (4sU: scSLAM-seq^18^, sci-fate^17^, NASC-seq^16^, scNT-seq^14^, Well-TEMP-seq^20^, and others^19, 71^) or 5-ethynyl-uridine (5EU: scEU-seq^15^, spinDrop^70^, TEMPOmap^21^, and others^72^).

Our study considers two types of labeling experiments: Pulse and chase^15^. Pulse experiments consist of labeling *n* cell cultures, starting at times *t_j_*, *j* ϵ {1, …, *n*}. Conversely, in chase experiments, cells are exposed to nucleoside analaogues long enough (*e.g.*, more than 24 hours), resulting in only labeled transcripts. Following, these labeled transcripts are washed out, starting at times *t_j_*. Similar to the pulse experiment, chase experiments include, in general, washing out at *n* different times. Finally, in both types of experiments, all cells are sequenced at a time *t_f_*, naturally defining the labeling time (or duration) by *τ*^(*j*)^_*l*_ = *t_f_* − *t_j_*.

Pulse and chase experiments allow measuring the production of mRNA. Here, we estimate cell-specific transcription and degradation rates, similar to a previous proposal in the scEU-seq study^15^. Specifically, for a particular gene, we assume mRNA levels to evolve according to

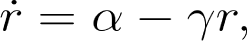

with transcription rate, and degradation rate. The corresponding solution is given by

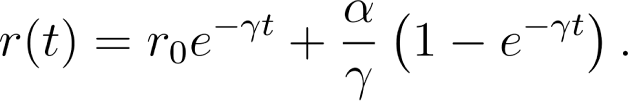

Note that here, we assume gene-specific models, *i.e.*, gene-gene interactions are neglected. In the following, we will identify mRNA measurements from pulse and chase experiments by the superscripts (*p*) and (*c*), respectively.

#### Pulse experiments

Pulse experiments study the production of labeled RNA. Since labeling starts at *t_k_*, and no labeled transcripts exist before, the abundance of labeled mRNA *r_l_* at times *t_k_* and *τ*^(*k*)^_*l*_ is given by

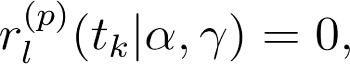

And

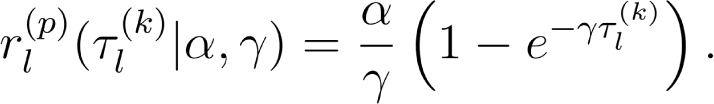

#### Chase experiments

In chase experiments, mRNA degradation is studied by washing out labeled transcripts. Thus, labeled mRNA *r*^(*c*)^_*l*_ at time τ^(*k*)^_*l*_ follows

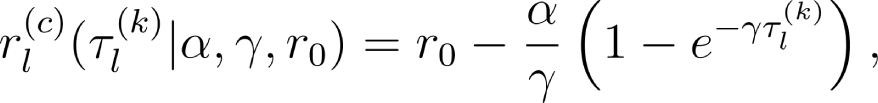

where corresponds to the mRNA level when starting to wash out labeled transcripts. Before washing out labeled mRNA, no unlabeled transcripts are present, and, thus, their abundance at time τ^(*k*)^_*l*_ is modeled as

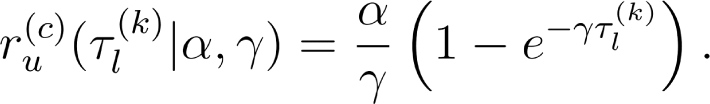

#### Parameter inference

Considering measurements from both chase and pulse experiments, we denote the respective set of cells by *C* and *P*. To estimate cell *j* and gene *g* specific model parameters α^(*j,g*)^, γ^(*j,g*)^, and *r*^(*j,g*)^_0_, we proceed as follows:

1. Consider cell *j* and its PCA representation *z*^(PCA)^_*j*_. For each labeling duration *k*, we determine the distance in PCA space between the reference cell *j* to each cell with labeling duration *τ*^(*k*)^_*l*_. For each gene *g*, we then identify the 20 nearest cells with non-trivial expression in *g*. These cells, as well as all closer neighbors (with zero counts), define the set *N*^(*k*)^_*g*_, which we consider for parameter inference.
2. To estimate model parameters, we minimize the quadratic loss *l* defined as

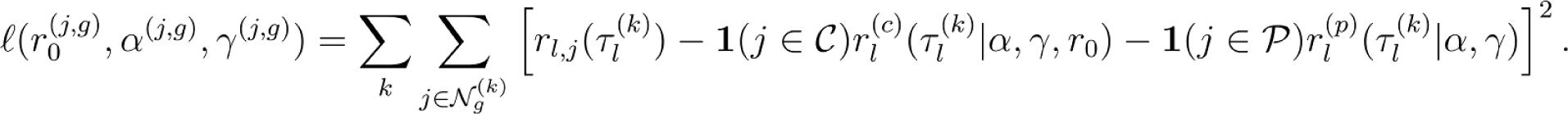

Here, **1**(*x* ϵ *Χ*) denotes the characteristic function taking equalling 1 if *x* ϵ Χ, and 0 otherwise. We note that estimating the parameters of a pulse (chase) experiment requires at least two (three) labeling durations.

Our approach differs from the scEU-seq study^15^ mainly in two ways: First, we base our analysis on total RNA, not spliced RNA. We reasoned that this approach circumvents limitations of identifying unspliced and spliced counts. Second, we infer rates for all genes and not only those changing significantly during development.

#### Method comparison

To benchmark the performance of different approaches, we identified and ranked potential drivers of every lineage, using each approach. We compared this ranking to a curated list of known lineage markers and regulators. If the literature-based gene set were complete, an optimal method would rank the corresponding genes highest. Consequently, for each method, we quantified its performance as follows: First, consider a lineage, a set of known drivers *D*, and a method *m*. Further, denote the set of genes by *G*, and for *g* ϵ *G*, identify its assigned rank by a superscript, *e.g.*, *g*^(*j*)^ for the *j*-th ranked gene *g*. Next, for each threshold *N* ϵ ℕ and *N* ≤ |*G*|, we computed how many known markers/regulators were ranked among the top *N* genes with

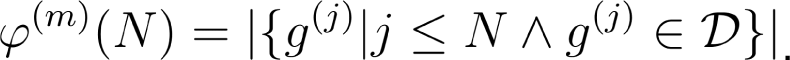

Thus, we call an assignment optimal when 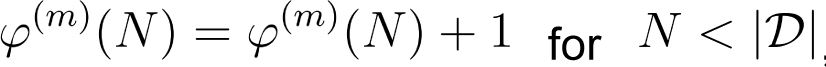, and 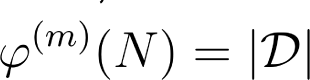 otherwise. Next, for each method *m*, we computed the area under the curve AUC(*m*) of φ^(*m*)^, *i.e.*,

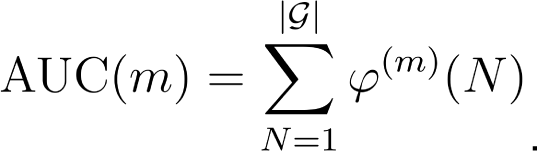

For each method, we then computed its relative area under the curve AUC_rel_(*m*) as

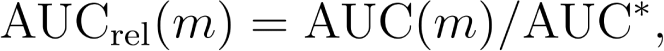

with

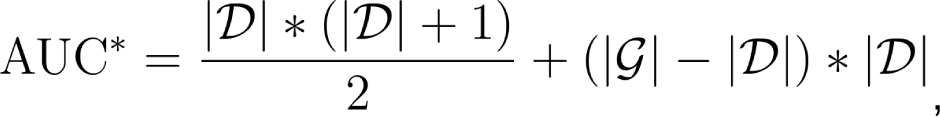

*i.e.*, the area under the curve of an optimal assignment.

### Datasets

Unless stated otherwise, all functions were run with default parameters. For Scanpy-based workflows^85^, we computed PCA embeddings, neighbor graphs, and UMAP embeddings^86, 87^ with the `scanpy.tl.pca`, `scanpy.pp.neighbors`, and `scanpy.tl.umap` functions, respectively.

For analyses based on CellRank 2 kernels, the kernel method `compute_transition_matrix` computed transition probabilities. We used the `GPCCA` estimator functions to compute macrostates (`compute_macrostates`)^27^, define terminal states (`set_terminal_states`), compute fate probabilities (`compute_fate_probabilities`), and identify putative lineage drivers (`compute_lineage_drivers`). To order the putative regulators according to their peak expression in pseudotime, we first fitted generalized additive models (GAMs) to describe gene expression change over pseudotime with `cellrank.models.GAM`. Following, we visualized the putative cascade of regulation with the `cellrank.pl.heatmap` function.

### Human hematopoiesis

All analyses were conducted on the dataset pre-processed by the original study^31^, and subsetted to the normoblast, dendritic, and monocyte lineages according to the provided cell type annotation (“HSC”, “MK/E progenitors”, “Proerythroblast”, “Erythroblast”, “Normoblast”, “cDC2”, “pDC”, “G/M prog”, “CD14+ Mono”).

#### Pseudotime-based analysis

After subsetting the data, we computed the nearest neighbor graph on the pre-computed MultiVI^85, 88^ latent space and the UMAP embedding with Scanpy. Following, we computed 15 diffusion components^89^ (scanpy.tl.diffmap) to then assign diffusion pseudotime values using Scanpy’s dpt function with n_dcs=6 (refs.^2, 89^). We identified the root cell as the hematopoietic stem cell with the largest fifth diffusion component.

We computed the transition matrix with CellRank 2’s PseudotimeKernel and thresholding_scheme=”soft”, and computed six macrostates^27^. We defined the terminal states “pDC”, “CD14+ Mono”, “Normoblast”, and “cDC2” that corresponded to the four macrostates with the largest macrostate purity. After quantifying fate probabilities, we identified putative pDC lineage drivers with our correlation-based procedure, restricted to the HSC and pDC clusters (lineages=[“pDC”] and clusters=[“HSC”, “pDC”]). We quantified the corresponding gene trends in the same way as described in our previous work^22^.

#### RNA velocity-based analysis

To infer RNA velocity, we generally followed the instructions provided by scVelo’s^12^ tutorials. First, we filtered for genes expressed in at least 20 cells in both unspliced and spliced counts with scVelo’s `scv.pp.filter_genes` function, and normalized counts with `scv.pp.normalize_per_cell`. The neighbor graph was again computed on the MultiVI latent space, followed by count imputation through first-order moments with scVelo’s `scv.pp.moments` function. We then inferred RNA velocity with the `scv.tl.recover_dymanics` function.

To quantify cellular fate, we computed transition matrices with the VelocityKernel and ConnectivityKernel, and combined them with 0.8 and 0.2 weight, respectively, as proposed by the CellRank 1 workflow. We computed macrostates and fate probabilities using the GPCCA estimator as described for the pseudotime-based analysis. As the RNA-velocity-based analysis did not identify the cDC cluster as a macrostate, we computed three macrostates corresponding to the terminal states normoblasts, monocytes, and pDCs.

### Embryoid body development

#### Data preprocessing

We followed Scanpy’s workflow to process the raw count matrix: As a first step, we filtered out genes expressed in less than 10 cells (`scanpy.pp.filter_genes` with `min_cells=10`). Following, we removed cells with more than 17.500 counts, cells for which more than 15% of counts originate from mitochondrial genes, and cells expressing more than 3500 genes. Following, we size normalized cells to 10.000 (`scanpy.pp.normalize_total` with `total_sum=1e4`), applied a log1p transformation (`scanpy.pp.log1p`), and annotated highly variable genes with `scanpy.pp.highly_variable_genes`. We based all further analyses on these highly variable genes and the marker genes identified by the study introducing the embryoid body development dataset^35^. The neighbor graph was computed for 30 neighbors using 30 principal components (PCs).

### CytoTRACEKernel analysis

To compute the CytoTRACE score^25^, we first imputed the normalized count matrix by first-order moments with scVelo’s `scvelo.pp.moments` function. The score itself was calculated with the `compute_cytotrace` method of the `CytoTRACEKernel`. We computed the transition matrix with the soft thresholding scheme (`thresholding_scheme=”soft”`) and `nu=5`. Putative drivers of the endoderm lineage were identified by focusing on the stem cell and endoderm clusters (`lineages=[“EN-1”]` and `clusters=[“ESC”]`).

#### Pseudotime construction

To compute DPT^2^, we calculated diffusion components (scanpy.tl.diffmap) and identified the putative root cell as the minimum in the first diffusion component. We then assigned DPT values using Scanpy’s dpt function.

For the Palantir pseudotime^7^, we used the corresponding Python package and followed the steps outlined in its documentation. As a first step, we computed the first five diffusion components with palantir.utils.run_diffusion_maps with n_components=5. Following, we identified the multi-scale space of the data (palantir.utils.determine_multiscale_space) and imputed the data using MAGIC^84^ (palantir.utils.run_magic_imputation). Finally, we computed the Palantir pseudotime via palantir.core.run_palantir using the same root cell as for our DPT analysis, and num_waypoints=500.

### Mouse embryonic fibroblast (MEF) reprogramming

#### Data preprocessing

For analyzing the dataset of mouse embryonic fibroblast (MEF) reprogramming towards induced pluripotent stem cells (iPSCs)^23^, we subsetted to the *serum* condition and added the category “MEF/other” to the cell set annotations. Following, we computed the PCA embedding and nearest neighbor graph.

#### WOT-based analysis

To construct transport maps, we used the wot package^23^ and followed the provided tutorials. First, we instantiated an optimal transport (OT) model (wot.ot.OTModel with day_field=”day”) and computed the transport maps next (compute_all_transport_maps). We defined the target cell sets based on the provided cell type annotation, and quantified WOT-based fates towards the last experimental time point through the OT model’s fates function with at_time=18.

#### *RealTimeKernel*-based analysis

For our *RealTimeKernel*-based analysis, we relied on the transport maps computed with wot. When constructing the transition matrix, we considered within-time-point transitions for every experimental time point and weighed them by 0.2 (self_transitions=”all”, conn_weight=0.2).

To construct the real-time-informed pseudotime, we symmetrized the global transition matrix and row normalized it. The symmetrized matrix defined the _transitions_sym attribute of Scanpy’s DPT class. Following, we computed diffusion components with the DPT class’ compute_eigen function. The root cell for DPT was identified as an extremum of the most immature cell state within the first experimental time point in diffusion space. Here, we selected the maximum in the first diffusion component. Finally, we computed DPT itself with scanpy.tl.dpt.

We computed fate probabilities towards the four terminal states according to our canonical pipeline, *i.e.*, identification of four macrostates followed by fate quantification.

#### Pseudotime construction

To assign each observation its DPT value, irrespective of experimental time points, we computed diffusion maps (scanpy.tl.diffmap) and identified the root cell as the maximum value in the first diffusion component. Following, we computed DPT with scanpy.tl.dpt.

For constructing the Palantir pseudotime, irrespective of experimental time points, we followed the same steps as described for the embryoid body development data.

### Pharyngeal endoderm development

#### Data preprocessing

The pharyngeal endoderm development dataset provided by the original study^40^ had already been filtered for high-quality cells and genes. Consequently, we directly quantified highly expressed genes using Scanpy’s^85^ `highly_variable_genes` function. Following, we computed the PCA embedding and nearest neighbor graph based on 30 PCs and 30 neighbors (`n_pcs=30`, `n_neighbors=30`).

#### *RealTimeKernel*-based analysis

To study the pharyngeal endoderm development dataset with the *RealTimeKernel*, we followed the `wot` tutorials to compute transport maps: First, we instantiated an OT model (`wot.ot.OTModel` with `day_field=”day”`) and computed the transport maps next (`compute_all_transport_maps`). For the *RealTimeKernel*, we considered within-time-point transitions for every experimental time point and weighed them by 0.1 (`self_transitions=”all”`, `conn_weight=0.1`).

We estimated terminal states using the *GPCCA* estimator with default settings by computing 13 macrostates and selecting the known terminal clusters. After calculating fate probabilities, for each lineage, we identified putative driver genes with `GPCCA.compute_lineage_drivers` by restricting the analysis to progenitors of the corresponding lineage and excluding cell cycle, mitochondrial, ribosomal, and hemoglobin genes^90^.

To study mTEC development, we subsetted to the early thymus, UBB, parathyroid, cTEC, and mTEC clusters and processed the data as described for the entire dataset. We computed the UMAP embedding using Scanpy’s `umap` function. To compute the transition matrix, we proceeded in the same manner as described for the entire dataset. For terminal state identification, fate quantification, and driver analysis, we followed the standard CellRank 2 pipeline.

#### WOT-based analysis

To identify putative drivers of the mTEC lineage with WOT, we used the same transport maps as in our *RealTimeKernel*-based analysis. We defined the target cell sets based on the provided cell type annotation considering only observations from the last time point and computed the pullback distribution from the mTEC cluster at embryonic day (E) 12.5 to E10.5 cells as it consists progenitor cells (pull_back). The sequence of ancestor distributions was quantified with the transport model’s trajectories method. WOT identifies putative drivers of the mTEC lineage as genes differentially expressed in cells most fated towards the mTEC cluster. We used the wot.tmap.diff_exp function to construct the corresponding gene ranking.

#### Classical differential expression analysis

As an alternative means to identify putative drivers of the mTEC lineage based on the fate probabilities assigned by our *RealTimeKernel*-based analysis, we defined two groups of cells within the general progenitor pool: those with mTEC fate probability greater than 0.5, and all other progenitor cells. We then identified differentially expressed genes between putative mTEC progenitors and all others with Scanpy’s rank_genes_groups function.

### Intestinal organoids

#### Data preprocessing

To preprocess the dataset of intestinal organoids, we first excluded DMSO control cells and cells labeled as tuft cells. Following, we removed genes with less than 50 counts, size normalized total and labeled counts, and identified the 2000 most highly variable genes with `scvelo.pp.filter_and_normalize`. The neighbor graph was constructed based on 30 PCs and 30 neighbors. Finally, we computed first-order-smoothed labeled and total mRNA counts.

#### Parameter estimation

To estimate kinetic rate parameters, we made use of our new inference scheme for metabolic labeling data implemented as part of the scVelo package. We first masked observations according to their labeling time with scvelo.inference.get_labeling_time_mask. Next, we computed pairwise distances between observations in PCA space, and sorted observations in ascending order for each time point using scvelo.inference.get_obs_dist_argsort. This information allowed us to identify, for each cell and gene, how many neighbors to consider during parameter estimation to include 20 non-zeros observations smoothed by first-order moments. This calculation was performed via scvelo.inference.get_n_neighbors. Finally, we estimated model parameters based on smoothed labeled counts with scvelo.inference.get_parameters.

#### Labeling velocity-based analysis

To quantify cell-specific fates, we first computed labeled velocities based on the estimated parameters. Following, we computed a transition matrix by combining the VelocityKernel and ConnectivityKernel with a 0.8 and 0.2 weight, respectively. We then inferred 12 macrostates and fate probabilities toward the known terminal states. Lineage-specific drivers were identified by restricting the correlation-based analysis to the corresponding terminal state and stem cell cluster. For putative driver gene ranking based on gene expression, we correlated fate probabilities with smoothed labeled counts.

#### Dynamo-based analysis

To analyze the intestinal organoid data with dynamo^24^, we followed the tutorials provided in the documentation of the Python package. As a first step, this required us to compute the ratio of new to total RNA with `dynamo.preprocessing.utils.calc_new_to_total_ratio` followed by first-order moment imputation of total and new RNA using `dynamo.tl.moments` with our connectivity matrix and `group=”time”`. Dynamo’s `dynamo.tl.dynamics` estimated the velocities with function arguments `model=”deterministic”`, `tkey=”time”`, and `assumption_mRNA=”ss”`.

Following velocity estimation, we quantified fixed points by following dynamo’s corresponding pipeline: First, we projected the high-dimensional velocity field of new RNA onto the UMAP embedding using `dynamo.tl.cell_velocities` with `ekey=”M_n”` and `vkey=”velocity_N”`. Fixed points were then identified by calling `dynamo.tl.VectorField` with `basis=”umap”` and `dynamo.vf.topography`. As a final step, we identified all stable fixed points.

Given the stable fixed points of the system, we identified putative genes regulating cell differentiation towards them using dynamo’s least action path analysis. As a first step, this workflow required us to compute a UMAP embedding based on new RNA with `dynamo.tl.reduceDimension` and `layer=”X_new”`, followed by the projection of the velocity field onto the PCA space (`dynamo.tl.cell_velocities` with `basis=”pca”`) and learning a vector field function based on this projection (`dynamo.tl.VectorField` with `basis=”pca”`). Next, we defined terminal states as the 30 nearest neighbors in UMAP space of each stable fixed point. For initial states, we computed the 30 nearest neighbors of unstable fixed points of the stem cell cluster. To compute the least action paths and account for uncertainty in initial and terminal state assignment, we randomly sampled ten pairs of initial and terminal cells and estimated the paths between them with `dynamo.pd.least_action`. Dynamo’s `dynamo.pd.GeneTrajectory` class then identified genes associated with the emergence of a terminal state. The pairwise sampling of initial and terminal cells defined the confidence bands of dynamo’s gene rankings shown in Fig. 3d.

#### RNA velocity-based analysis

Conventional RNA velocity was estimated with scVelo’s dynamical model by running `scvelo.tl.recover_dynamics`. To execute this function, we first preprocessed the raw data with `scvelo.pp.filter_and_normalize` to remove genes expressed in less than 50 cells (`min_counts=50`), size normalizing spliced and unspliced counts, and subsetting to the 2000 most highly variable genes (`n_top_genes=2000`). Following, we computed the PCA embedding, calculated the neighbor graph with 30 PCs and 30 neighbors, and smoothed unspliced and spliced counts by first-order moments.

We combined the VelocityKernel and ConnectivityKernel weighted by 0.8 and 0.2, respectively, to estimate the cell-cell transition matrix. Next, we identified terminal states and corresponding fates and putative driver rankings following the canonical CellRank 2 pipeline.

## Notes

https://github.com/theislab/cellrank

https://github.com/theislab/cellrank2_reproducibility

