## Supplementary Notes for "Unified fate mapping in multiview single-cell data"

### Supplementary Note 1 - Limitations of RNA Velocity

Although RNA velocity has been tremendously successful since its introduction[1], several recent publications[2–8] highlight the limitations of traditional RNA velocity models (velocyto[1] and scVelo[9]). These shortcomings include

- **Gene structure bias:** RNA velocity is a proxy for the current state of transcriptional regulation fundamentally based on faithfully recovering both spliced and unspliced counts from the same cell. However, recovering sufficient unspliced transcripts across many genes is difficult in practice since polyadenylation and splicing happen mostly simultaneously[10]. In fact, the majority of captured unspliced counts result from internal priming events where the poly(T) primers bind to poly(A) stretches in intronic regions of the unspliced transcript[1,6,11]. Consequently, unspliced counts of a certain gene can, for example, depend on
    - the length of the gene[6],
    - the number, length, and position of intronic regions within the gene[2],
    - the amount of poly(A) stretches within introns, or
    - the relative position of poly(A) stretches within an intron.

  Each case results in biases that are difficult to identify and impossible to control externally. Specifically, if key driver genes for a biological process lack sufficient unspliced counts for any of these reasons, the corresponding velocity estimates will most likely differ significantly from the ground truth. In this case, genes with large numbers of unspliced reads dominate the overall velocity of cells. Contrastingly, for spliced transcripts, the poly(T) primers bind to the poly(A) tail.
- **State and time-dependent reaction parameters:** RNA velocity is based on a model of the mRNA lifecycle in which transcripts are created at rate $\alpha$, spliced at rate $\beta$, and degraded at rate $\gamma$. Traditional velocity models assume gene-specific constant rates during induction and repression, respectively. However, practical examples, including some applications from the original publications[1,9], have shown that both splicing and degradation rates vary across cell types and states[2,12]. Further, transcription rates vary over time, for example, in the case of transcriptional bursting events during erythroid maturation[5,13]. Such time and state-dependent kinetic parameters violate assumptions of traditional RNA velocity models leading to velocity trends inconsistent with prior biological knowledge, for example, in hematopioeisis[2,5,13,14].
- **Gene-gene interactions:** Current velocity methods[1,9] fit rate parameters in a gene-specific fashion. This assumption ignores the fundamentals of gene-gene regulation.
- **Batch effects:** RNA velocity is based on the ratio of spliced to unspliced counts. This ratio is not preserved across batches or data integration approaches. Thus, spliced and

unspliced layers cannot be processed independently, making it difficult to infer RNA velocity for frequently generated atlas-scale datasets[15,16].

- **Noisy, stochastic expression kinetics:** State-of-the-art single-cell sequencing protocols yield noisy and sparse mRNA transcript counts[17,18]. This aspect holds in particular for unspliced transcripts, which are harder to detect than their spliced counterparts. Currently, most RNA velocity approaches handle the high noise level by smoothing gene expression before inference[1,9], thereby ignoring the stochastic nature of gene expression.
- **Time scales of splicing:** Splicing kinetics determine the time delay between unspliced and spliced transcripts. However, this temporal delay cannot be controlled externally. Therefore, the time horizon over which RNA velocity extrapolates cellular states is fixed and may be incompatible with typical time scales in a biological system of interest.
- **Parameter non-identifiability:** RNA velocity is traditionally inferred based on a two-dimensional system of ordinary differential equations (ODEs). In the steady-state setting proposed by velocyto[1], the first method estimating RNA velocity, gene-specific parameters could only be estimated up to a multiplicative constant and are thus not comparable across genes. The scVelo[9] approach partially addressed this issue through dynamical modeling and post-hoc estimation of a gene-shared latent time. While scVelo's inference scheme rendered kinetic parameters comparable across genes it left a global multiplicative constant that precludes estimation of kinetic rates in their native units.

Several recently proposed models attempt to overcome some of these limitations and can be grouped as follows:

- **Model extension towards other modalities:** While the original RNA velocity model is based on the time delay between spliced and unspliced mRNA processing stages, emerging models include other molecular layers, including
    - mRNA and surface proteins[19],
    - mRNA and chromatin accessibility[20–22],
    - open and closed chromatin[23],
    - new and total transcripts in single-cell metabolic labeling assays[3,24,25].

  By including other modalities, these approaches relax the dependency on gene structure and try to estimate kinetic rate parameters more accurately through more inclusive models.
- **Stochastic models of splicing kinetics:** While both scVelo[9] and velocyto[1] model average smoothed expression kinetics, recent approaches[7,26] formulate stochastic extensions to account for high noise levels in unspliced transcript abundance. However, these approaches are theoretical frameworks with no practical implementation.
- **Adapted velocity models:** scVelo and velocyto solve a system of two coupled ODEs per gene. More recent approaches formulate different models for extracting directional information from the spliced/unspliced-count ratio. Approaches include radial basis function fitting[27], a density-adaptive model to overcome parameter non-identifiability[8],

and graph-convolutional neural networks to estimate state and time-dependent reaction parameters[2,5,28].

Although these approaches address some of the velocity issues we outlined above and may be used directly through CellRank 2's *VelocityKernel,* they often do so at the cost of introducing new heuristics unlikely to generalize across datasets. These heuristics include estimation based on scVelo's steady-state model[28] or assuming that time differences between two cell states are inversely proportional to the density of sampled cells that lie between them[8].
