## Supplementary Tables for "Unified fate mapping in multiview single-cell data"

### Supplementary Table 1

Distribution of cells across experimental time points in mouse embryonic fibroblast dataset.

| Day | 0 | 0.5 | 1 | 1.5 | 2 | 2.5 | 3 | 3.5 |
|---|---|---|---|---|---|---|---|---|
| Number of cells | 4556 | 3449 | 3648 | 1956 | 6981 | 6734 | 6777 | 7355 |

| Day | 4 | 4.5 | 5 | 5.5 | 6 | 6.5 | 7 | 7.5 |
|---|---|---|---|---|---|---|---|---|
| Number of cells | 8962 | 7127 | 7227 | 6550 | 8422 | 3111 | 6507 | 5061 |

| Day | 8 | 8.25 | 8.5 | 8.75 | 9 | 9.5 | 10 | 10.5 |
|---|---|---|---|---|---|---|---|---|
| Number of cells | 3815 | 3829 | 3573 | 3088 | 2982 | 2266 | 2051 | 1941 |

| Day | 11 | 11.5 | 12 | 12.5 | 13 | 13.5 | 14 | 14.5 |
|---|---|---|---|---|---|---|---|---|
| Number of cells | 2238 | 2164 | 2429 | 2253 | 2145 | 2034 | 3758 | 2723 |

| Day | 15 | 15.5 | 16 | 16.5 | 17 | 17.5 | 18 |
|---|---|---|---|---|---|---|---|
| Number of cells | 3717 | 4851 | 3422 | 4645 | 3678 | 4068 | 3799 |
